# *Babesia divergens* egress from host cells is orchestrated by essential and druggable kinases and proteases

**DOI:** 10.1101/2022.02.17.480550

**Authors:** Brendan Elsworth, Caroline D. Keroack, Yasaman Rezvani, Jacob A. Tennessen, Samantha A. Sack, Aditya S. Paul, Cristina K. Moreira, Marc-Jan Gubbels, Kourosh Zarringhalam, Manoj T. Duraisingh

## Abstract

A unique aspect of apicomplexan biology is the requirement for egress from and invasion into host red blood cells (RBCs). The cellular mechanisms and molecular mediators of RBC egress and invasion remain poorly characterized in *Babesia* spp., a group of parasites of veterinary importance and emerging cause of zoonotic disease. Through the use of video microscopy, transcriptomics, and chemical genetics we have implicated signaling, proteases and gliding motility in egress and/or invasion by *Babesia divergens*. We developed CRISPR/Cas9 and two inducible knockdown systems to perform a genetic screen of putative mediators of egress. We found that proteases ASP2 and ASP3 are required for invasion, and the latter is also required for egress. Strikingly, parasites continue to replicate intracellularly in the absence of the protein kinases, PKG or CDPK4, indicating that they are required for exit from the replication cycle and egress. These essential molecules present druggable targets for *Babesia spp*. All together we have established a molecular framework for the spread of infection through host RBCs, with egress of *B. divergens* more closely resembling *T. gondii* than the more evolutionarily related *Plasmodium* spp.

**Highlights:**

- Egress in *Babesia divergens* requires host cell lysis and parasite motility
- Transcriptomics can be used to identify egress and invasion proteins
- Knockdown of the proteases, ASP2 and ASP3, inhibit egress and invasion
- Inhibition of PKG or CDPK4 signaling results in continued intracellular replication

## Introduction

*Babesia* spp. are important tick-borne pathogens that infect, grow in, and eventually destroy the red blood cells (RBCs) of their vertebrate host. Several *Babesia* spp., including *B. microti*, *B. divergens* and *B. duncani*, are emerging zoonotic pathogens and can cause severe and even fatal disease. *Babesia* spp. also cause significant economic losses globally due to disease in livestock and companion animals, especially *B. divergens*, *B. bovis* and *B. bigemina* in cattle (Bock et al., 2004). Current treatment and prevention options for human and veterinary babesiosis are limited and suffer from poor efficacy, spontaneous resistance, severe side effects or render the animal products unsuitable for human consumption.

The *Babesia* genus is within the Apicomplexa phylum, which consists of single-celled obligate intracellular parasites that cause a significant burden on human and animal health. Apicomplexa include the human and veterinary pathogens, *Babesia* spp., *Plasmodium* spp., *Toxoplasma* spp., *Cryptosporidium* spp., *Theileria* spp. and *Eimeria* spp., amongst others. *Babesia* spp. are most closely related to *Theileria* spp. and *Plasmodium* spp., which all share the same vertebrate RBC niche for part of their lifecycle. The vast majority of apicomplexan cellular and molecular biology research has been performed in *Plasmodium* spp. and *Toxoplasma gondii*. Knowledge of a wider range of apicomplexan biology will hone in on conserved and essential functions that could be key targets to develop broad spectrum anti-apicomplexan drugs. There are many similarities between *Plasmodium* spp., *Toxoplasma* spp. and *Babesia* spp., such as the presence of an apicoplast and apical invasion organelles. However, there are notable differences, such as *Babesia* spp. degrading their parasitophorous vacuole immediately after invasion and replicating in the host cell cytoplasm. The mechanisms of cell division also vary between apicomplexan parasites (reviewed in Gubbels et al. (2020)). In this study, a replication cycle refers to one cell undergoing division to form two (*Babesia* spp. and *T. gondii*) or more (*Plasmodium* spp.) daughter cells. A lytic cycle refers to the time from when a host cell is first invaded to when parasites egress from the same host cell, which in the case of *Babesia* spp. and *T. gondii* may include multiple replication cycles (Figure S1A). Within the lytic cycle of *B. divergens*, the parasite undergoes one (∼66% per invasion event), two (∼33% per invasion event) and rarely three or more (∼1% per invasion event) replication cycles to produce two, four and eight or more daughter parasites, respectively, before egressing (Figures S1A and S3H) (Cursino-Santos et al., 2016). *T. gondii* also produce two daughter cells from one replication cycle but within a single lytic cycle *T. gondii* can produce hundreds of daughter cells. In contrast, the lytic cycle of *Plasmodium* spp., also referred to as the intraerythrocytic developmental cycle (IDC), always contains a single replication cycle. *Plasmodium* spp. asexual stages divide by schizogony in which the nucleus replicates multiple times before a final round of nuclear division that coincides with cytokinesis and a single replication cycle can form 16-32 daughter cells for *P. falciparum* asexual stages.

Significant effort has been utilized to develop inhibitors of egress in *Plasmodium* spp. and *T. gondii* (reviewed in (Caldas and De Souza, 2018; Singh and Chitnis, 2017). Egress remains poorly characterized in *Babesia* spp. Many of the molecular players in the host cell egress program are shared between *Plasmodium* spp. and *T. gondii*, including cGMP-dependent kinase (PKG), calcium-dependent protein kinases (CDPKs), aspartyl proteases, lipid and calcium signaling (Bisio and Soldati-Favre, 2019; Tan and Blackman, 2021). However, there are key differences, including the role of protein kinase A (PKA) to repress egress in *T. gondii,* whereas PKA is instead required for invasion in *P. falciparum* (Jia et al., 2017; Uboldi et al., 2018; Wilde et al., 2019). Egress in these organisms begins when either intrinsic or extrinsic signals activate signaling pathways in the parasite. Activation of the signaling pathways ultimately result in the release of parasite secretory organelles, the micronemes/exonemes, into the host cell cytoplasm. The contents of these organelles include proteases, phospholipases and perforin-like proteins (PLPs), which help permeabilize and eventually rupture the parasitophorous vacuole membrane (PVM) and host cell membrane, allowing the parasite to egress from the host cell.

Here, we used video microscopy, transcriptomic and chemical genetic methods to produce a foundational framework defining the key cellular features of *B. divergens* egress and invasion as well as identify the molecular mediators of these processes through functional analysis. We have developed stable transfection, a CRISPR/Cas9 system and inducible knockdown systems for *B. divergens*. These systems were employed to determine that the *B. divergens* kinases, PKG and CDPK4, and the aspartyl proteases, ASP2 and ASP3, are required for separate sequential steps in egress and invasion of the host RBC. Strikingly, inhibition of egress signaling resulted in continued intracellular replication within a single RBC, suggesting the default pathway in these parasites is to continue replicating in the absence of an egress signal.

## Results

### cGMP analogs and phosphodiesterase inhibitors efficiently induce egress of *B. divergens*

Like *T. gondii*, *B. divergens* replication is difficult to synchronize and egress does not always occur at the end of a replication cycle. Chemically induced egress has proven powerful in studying the process in *T. gondii*. To make the study of egress accessible in *B. divergens*, we developed a method to induce egress. A flow cytometry-based assay was used to screen known *T. go* uction in *B. divergens* (Figures 1A, S1B and S1D). 8-Bromoguanosine 3’,5’-cyclic monophosphate (8-Br-cGMP), BIPPO and to a lesser extent, H89, induced egress (Figures 1A and S1D). All three compounds inhibited parasite growth with an IC_50_ between 1.3-11.2 µM (Figure S1F). BIPPO inhibits phosphodiesterases (PDE) which hydrolyze cGMP and cAMP in *P. falciparum* (Howard et al., 2015), whereas 8-Br-cGMP is a cell-permeable and hydrolysis-resistant analog of cGMP. Both compounds presumably lead to the activation of PKG, a well-characterized mediator of egress in *P. falciparum* and *T. gondii*. 8-Br-cGMP was used as the inducing agent for all further experiments. The rate and total number of parasites that egress is dependent on 8-Br-cGMP concentrations (Figure S1H). PKA activity is required to suppress egress in *T. gondii* but is conversely required for invasion in *P. falciparum* (Jia et al., 2017; Patel et al., 2019; Uboldi et al., 2018; Wilde et al., 2019). The PKA inhibitor, H89, strongly inhibited invasion of isolated *B. divergens* merozoites and weakly induced egress, suggesting that BdPKA may have dual roles in the parasite or the two PKAc orthologs may have separate roles in egress and invasion (Figures S1D and G). A summary of the putative *B. divergens* egress signaling pathway can be found in S1C.

**Figure 1.**
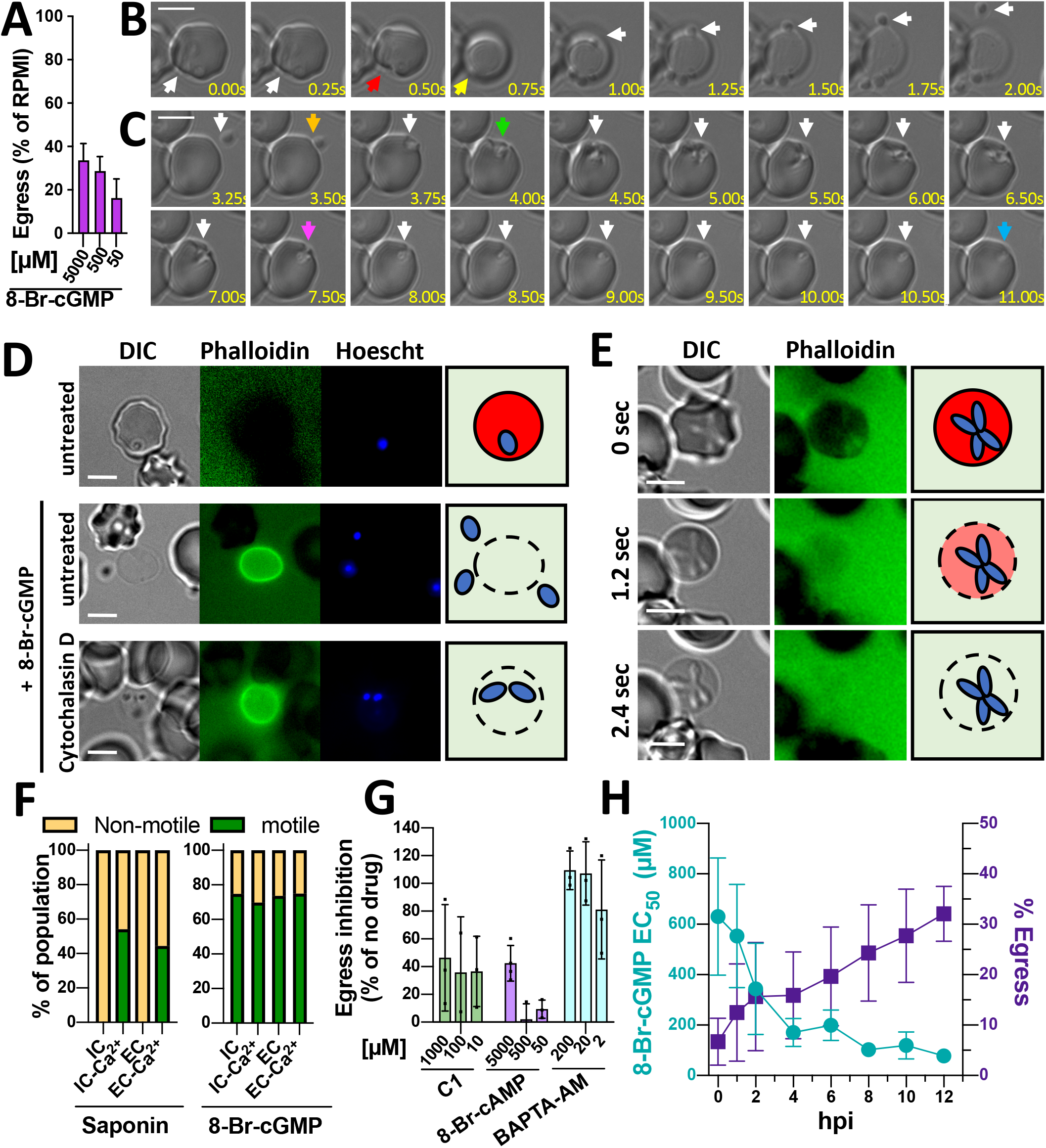
Egress and invasion by *B. divergens*. **(A)** Percentage of parasites that egress when treated with 8-Br-cGMP. Data is normalized to 0% being an RPMI treated control (B and C) Time-lapse microscopy of *B. divergens* egress **(B)** and invasion **(C)**. A and B follow the same parasite egressing from one cell and invading a new RBC. Arrows indicate as follows: Red - initial deformation of the RBC in egress. Yellow - ‘rounding-up’ associated with permeabilization of the RBC. Orange - initial RBC contact in invasion. Green - initial deformation of the RBC. Purple – beginning of internalization. Blue – completion of invasion. White – follows the parasite. **(D)** *B. divergens* iRBCs pre-treated with cytochalasin D and induced to egress with 8-Br-cGMP. Phalloidin stains the permeabilized RBC. **(E)** Time-lapse images showing RBC permeabilization by cytochalasin D*-*treated *B. divergens* during induced egress. Phalloidin is excluded from the intact RBC. **(F)** Percentage of parasites that become motile when the iRBC is permeabilized by saponin or induced to egress with 8-Br-cGMP in the stated buffer. IC and EC refer to intracellular and extracellular buffer with or without 2 mM calcium (Ca^2+^). Data is pooled from multiple experiments. >30 iRBCs were used for each condition. **(G)** Screen of small molecules for inhibition of 8-Br-cGMP mediated induced egress. Data is normalized to 100% as a no 8-Br-cGMP control and 0% being RPMI + 8-Br-cGMP. **(H)** Egress was induced throughout one replication cycle using a range of 8-Br-cGMP concentrations. The EC_50_ of 8-Br-cGMP is shown in teal and the percentage of parasites that egress at 2 mM 8-Br-cGMP is shown in purple. Data is normalized to 0% being a no 8-Br-cGMP treated control. For G and H, the mean ± SD of three biological experiments performed in technical triplicate is shown. The scale bar in A and B is 5 µM.

### The egress and invasion processes of *B. divergens*

*T. gondii* secreted lytic factors (e.g. PLP1) and host calpains damage the host cell, allowing the motile parasite to escape the permeabilized cell (Chandramohanadas et al., 2009; Kafsack et al., 2009; Moudy et al., 2001). In contrast, *P. falciparum* secreted proteases degrade the RBC cytoskeleton, leading to the to rupture of the RBC to release free merozoites, thus bypassing the requirement for gliding motility in egress (Das et al., 2017; Perrin et al., 2018; Thomas et al., 2018). We used video microscopy to follow 8-Br-cGMP mediated induced egress to better understand the mechanics of egress, motility and invasion in *B. divergens*. As has previously been observed, *B. divergens* egress is initiated when the intracellular parasite contacts and deforms the RBC (Figure 1B, red arrow)(González et al., 2019). Typically, within <0.5 s from initial deformation the RBC ‘rounds up’, a phenotype that has been observed in *P. falciparum*, which is thought to be related to permeabilization of the PVM and/or reorganization of the merozoites, however, the mechanism remains unclear, and is likely due to permeabilization of the RBC in *B. divergens* (Glushakova et al., 2018; Hale et al., 2017; Thomas et al., 2018). *B. divergens* parasites then become motile to escape the permeabilized RBC and search for new RBCs to invade. Upon contact with a new RBC, the parasite dramatically deforms the host cell around itself (Figure 1C, green to purple arrows), a process known as pre-invasion in *P. falciparum*, followed by the internalization phase when the parasite enters the RBC with relatively little deformation of the RBC (Figure 1C, purple to blue arrows, Video 1) (Gilson and Crabb, 2009). In some instances, the parasite becomes motile within the intact RBC before the RBC ruptures (Video 2).

### Motility is required for *B. divergens* merozoites to egress from permeabilized RBCs but not for permeabilization

The initial deformation of the RBC in egress appears to be caused by direct contact with the parasite, suggesting the parasites actinomyosin motor may be used to physically disrupt the host membrane (Figure 1B) (González et al., 2019). To test if motility is required during egress, 8-Br-cGMP was used to induce egress in the presence of phalloidin-Alexa Fluor 488, which selectively stains the cytoskeletons of permeabilized RBCs, and cytochalasin D, which inhibits actin polymerization and gliding motility. Cytochalasin D-treated parasites can be induced to lyse the host cell, but do not escape the permeabilized cell (Figure 1D and E, Video S3). Immediately preceding egress, the host cell becomes ‘ruffled’, but no localized deformation is observed, followed by the RBC rapidly showing a reduced diameter, becoming round and being infiltrated by phalloidin demonstrating the RBC membrane has been permeabilized (Figure 1E, Video S3). In contrast to *P. falciparum* which fractures the RBC cytoskeleton during egress (Figure S1J) (Glushakova et al., 2010), the permeabilized RBC remains relatively intact throughout egress in *B. divergens* (Figures 1B-E) (González et al., 2019). Together this data suggests that *B. divergens* secretes lytic factors to permeabilize the host cell and does not strictly require motility for host cell lysis, however, we cannot rule out that motility can contribute towards cell lysis as has been observed in *T. gondii* (Kafsack et al., 2009).

### Loss of host cell integrity induces *B. divergens* motility and egress

*T. gondii* can sense exposure to the extracellular environment through serum albumin, a drop in potassium and an increase in extracellular calcium, which induces microneme secretion and motility (Black et al., 2000; Brown et al., 2016; Moudy et al., 2001; Vella et al., 2021). *P. falciparum* can sense a drop in potassium, which enhances microneme secretion, however, egress and invasion efficiency is not dependent on the drop in potassium, making its biological role unclear (Pillai et al., 2013; Singh et al., 2010). *P. falciparum* also requires extracellular calcium for efficient invasion but it is not required for egress, microneme secretion or activation of the actino-myosin motor (Singh et al., 2010; Wasserman et al., 1982; Weiss et al., 2015b). To test if *B. divergens* motility is induced by exposure to extracellular conditions, we observed parasites after the RBC was lysed using a low concentration of saponin in buffers mimicking either intracellular (IC, 140 mM K^+^, 5 mM Na^+^) or extracellular (EC, 5 mM K^+^s, 140 Na^+^) concentrations of potassium and sodium, plus or minus calcium (2 mM). Upon saponin induced RBC lysis, parasites in EC-Ca^2+^ buffer became motile and were able to escape from the permeabilized RBC to a similar extent (Figure 1F, Video S4). In contrast, parasites in either IC or EC buffer without Ca^2+^ did not become motile (Figure 1F, Video S5). PKG activation, through the addition of 8-Br-cGMP, was able to bypass the requirement for extracellular calcium in either buffer to initiate gliding motility (Figure 1F). Together these results suggest that *B. divergens* motility is induced by exposure to extracellular concentrations of Ca^2+^, likely acting upstream of PKG, but does not respond to Na^+^ or K^+^ concentrations.

### Chemical inhibition of PKG, proteases and calcium release impair egress

To identify the molecular processes required for *B. divergens* egress, we used the flow cytometry-based egress assay to screen for compounds that inhibit 8-Br-cGMP-induced egress. The compounds were selected based on previous studies that have demonstrated egress inhibition in *P. falciparum* or *T. gondii*. BAPTA-AM, which chelates intracellular calcium, and Compound 1 (C1), an inhibitor of apicomplexan PKG, both inhibited 8-Br-cGMP induced egress (Figure 1G). Inhibitors of serine proteases, including TPCK, TLCK and PMSF, reduced induced egress at similar concentrations to those that inhibit *P. falciparum* gametocyte egress (Figure S1E) (Rupp et al., 2008). Inhibitors of other protease classes, which have been shown to inhibit *Plasmodium* spp. egress, produced only modest inhibition, including pepstatin, chymostatin and E64 (Figure S1E) (Lyon and Haynes, 1986; Salmon et al., 2001). Inhibitors of the lipid signaling pathway that inhibit egress in *T. gondii* and/or *P. falciparum*, including U73122 and propranolol which target PI-PLC and phosphatidic acid phosphatase, respectively, did not influence egress by *B. divergens* (Figure S1E) (Agarwal et al., 2013; Bullen et al., 2016; Moudy et al., 2001). The diacylglycerol kinase inhibitor R59022, an inhibitor of egress in *T. gondii* and egress to invasion in *P. falciparum*, instead led to enhanced egress in the presence of 8-Br-cGMP (Bullen et al., 2016; Paul et al., 2020). We note that concentrations of R59022 >20 µM resulted in visible RBC lysis and low-level lysis may occur at 20 µM, leading to a false egress signal in the flow cytometry assay (data not shown). In line with our flow cytometry assay, E64d treated parasites showed no obvious defect in egress, motility or invasion by video microscopy (Figure S1K). No egress events or permeabilized RBCs were observed when parasites were treated with TPCK or BAPTA-AM (Figure S1K, data not shown). Taken together, these data demonstrate a requirement for cGMP/PKG signaling, calcium signaling (both intracellular and extracellular), serine proteases and gliding motility for efficient egress by *B. divergens*.

### B. divergens can be induced to egress throughout its intracellular replication cycle

*T. gondii* can be induced to egress throughout the replication cycle, albeit at reduced efficiency during S and M/C phase (Gaji et al., 2011), whereas *P. falciparum* egress is restricted to a narrow window at the end of the lytic cycle (Collins et al., 2013; Howard et al., 2015). To test the ability of *B. divergens* to egress throughout replication, parasites were synchronized to a 20 minute window and egress was induced every 1-2 hrs. Immediately after invasion, fewer parasites egress at the highest concentration of 8-Br-cGMP (6.5% with 2 mM at 0 hpi) and higher concentrations are required to induce egress (EC_50_ 630 µM) (Figure 1H). As the parasite matures, the EC_50_ of 8-Br-cGMP required to induce egress decreases and the number of parasites that egress at the highest concentration increases (EC_50_ 78 µM and 32% egress with 2 mM at 12 hpi) (Figure 1H). Parasites that do not egress become pyknotic by the end of the 1 hr assay (Figure S1I). These results suggest that *B. divergens* can egress throughout the replication cycle, although it is more strongly primed to egress when parasites are fully mature.

### Identification of putative egress, motility and invasion genes through transcriptomics

The lytic cycle of *B. divergens* has been morphologically defined but remains poorly characterized at the molecular level in any *Babesia* spp. (Cursino-Santos et al., 2016). There are several *Babesia* spp. transcriptomes at different stages of the life cycle, however, there are no synchronous time course transcriptomes (Pedroni et al., 2013; Peloakgosi-Shikwambani, 2018; Rossouw et al., 2015; Silva et al., 2016; Ueti et al., 2020). We generated a synchronous transcriptome of one replication cycle (0-12 hrs) for *B. divergens*. Analysis of the transcriptome revealed 3990 of 4125 (96.73%) genes can be detected in *in vitro* culture, similar to previous studies in *P. falciparum* (∼90%) (Chappell et al., 2020; Otto et al., 2010) and *T. gondii* (∼85%) (Hassan et al., 2017; Hehl et al., 2015) (Figure 2A). 2190 (53.10%) genes showed expression changes over time of >1.5-fold. As a parallel strategy, a single-cell transcriptome was generated from an asynchronous culture. 3620 (87.76%) genes were detected, of which 560 (13.57%) display significant changes over time (≥2-fold-change, adj. p-value 0.001) (Figure 2A). To generate expression profiles from the single-cell data, a pseudo-time analysis was performed and the start of gene expression was identified by cross-correlation analysis between the bulk and single-cell expression curves (see methods and (Rezvani et al., 2022) for details). Genes that showed expression changes over time display a transcriptional cascade associated with “just-in-time” gene expression that has been observed in *Plasmodium* spp. and *T. gondii* (Figures 2B and C) (Behnke et al., 2010; Bozdech et al., 2003). The time of peak expression between the bulk and single-cell transcriptomes was well correlated for the majority of genes (Figure 2D).

**Figure 2.**
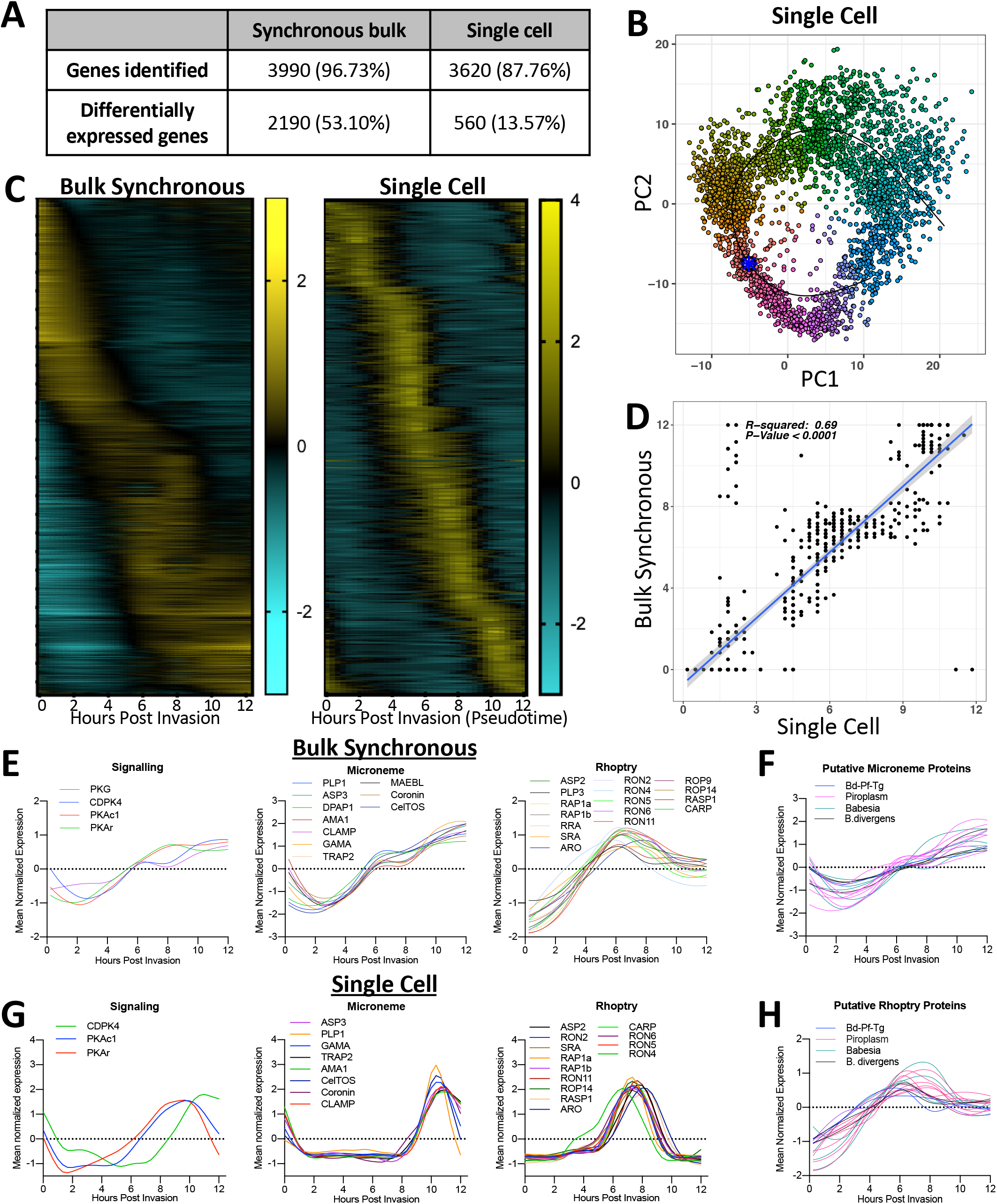
The *B. divergens* transcriptome reveals putative egress and invasion genes. **(A)** Expression profile of all differentially expressed genes. **(B)** PCA plot of the single-cell transcriptomics. **(C)** Number of genes identified by both transcriptomic approaches. **(D)** Correlation of timing of peak expression between both approaches. **(E-H)** The expression profiles from bulk synchronous and single-cell data of orthologs of known egress/invasion genes (E,G) or putative novel egress/invasion genes **(F,H)**.

93 *B. divergens* genes were identified as orthologs, or family members of, known egress, motility or invasion genes in apicomplexan parasites, 91 of which are expressed and 67 showed significant expression changes over time (Table S1). Genes that are expected to co-localize within the same subcellular compartment (e.g. micronemes) based on orthology, display similar expression profiles, which also occurs in *Plasmodium* spp. and *T. gondii* (Figures 2E, G and S2C) (Behnke et al., 2010; Besteiro et al., 2009; Young et al., 2008). Second messenger-responsive proteins, including PKG, PKAc1, PKAr, and CDPK4 remain high until 12 hpi when parasites naturally egress (Figures 2E and G). Other putative members of the egress signaling pathway did not display stage-specific expression indicative of a role in egress or invasion (Table S1). The cysteine and aspartyl proteases, DPAP1 (no direct ortholog) and ASP3 (ortholog of PfPMX/TgASP3), respectively, display expression profiles matching that of microneme proteins (Figures 2E and G). The aspartyl protease ASP2 (closely related to BdASP3) displays an expression profile matching rhoptry proteins (Figures 2E and G). Both BdASP2 and BdASP3 have higher sequence identity to PfPMX than to PfPMIX, however, subcellular localization based on their transcriptomic profiles suggest BdASP2 and ASP3 are functionally orthologous to PfPMIX and PfPMX, respectively. All nine PLP proteins in *B. divergens*, the orthologs of which are required for egress in *T. gondii* or *P. falciparum* sexual stages, are expressed in *in vitro* culture (Deligianni et al., 2013; Kafsack et al., 2009; Paoletta et al., 2021). Only PLP1 (TgPLP1/PfPLP3) and PLP3 (no direct ortholog) display an expression profile matching other microneme and rhoptry proteins, respectively, suggestive of a role in egress and invasion or post-invasion PV breakdown, respectively (Figures 2E and G).

Multiple invasion ligands were identified by orthology to known apicomplexan ligands and display expression profiles matching that of rhoptry or microneme proteins (Figures 2E and G). These included TRAP2/P18, RAP1 and AMA1, which all have a demonstrated role in *Babesia* spp. invasion (Figure 2E and G)(Terkawi et al., 2013; Wright et al., 1992; Zhou et al., 2006). The *B. divergens* orthologs of the *Plasmodium* spp. or *T. gondii* rhoptry protein, SRA, and microneme proteins, CLAMP, GAMA, MAEBL and CelTOS represent novel vaccine targets that have not been investigated in *Babesia* spp. (Amlabu et al., 2018; Arumugam et al., 2011; Bergmann-Leitner et al., 2010; Ghai et al., 2002; Sidik et al., 2016).

A comparative transcriptomic approach was used to identify novel proteins putatively involved in egress, motility or invasion. The Pearson correlation was calculated between the transcriptional profile of *B. divergens* (bulk RNAseq), *P. falciparum* and *T. gondii* orthologs, normalized to the length of their replication cycle, using existing datasets (Behnke et al., 2010; Otto et al., 2010). Comparing positively correlated genes across the genome between species pairs by timing of peak expression or by gene essentiality revealed only weak trends (Figures S2B, E-G). Three approaches were used to identify novel genes involved in egress or invasion: 1) identification of genes containing a signal peptide with an expression profile matching those of other microneme and rhoptry genes in *B. divergens*, 2) identification of orthologs that share late-stage expression between *B. divergens* and *P. falciparum* or *T. gondii*, and 3) identification of *B. divergens* orthologs of proteins localized to the rhoptries or micronemes by proteomics in *T. gondii* or *P. falciparum* (Dunkley et al., 2004; Lal et al., 2009; Swearingen et al., 2016). 104 genes were identified, 31 of which were identified by multiple methods and whose orthologs have no known function (Figures 2F and 2H; Table S2). Of the 8 genes found in all three species, 5 have a growth phenotype in the *T. gondii* CRISPR screen and the *P. falciparum piggyBac* screen (Table S2). Only one gene, Bdiv_039160, contains a known domain (TMEM121), which has no known function.

### A genetic screen of high priority candidates reveals PKG, CDPK4, ASP2 and ASP3 are essential for parasite proliferation

We focused on 11 high priority candidates based on orthology of apixomplexan egress genes, including the kinases, PKG, PKAc1, PKAc2, CDPK4, CDPK5 and CDPK7, the perforin-like proteins, PLP1 and PLP3, and the proteases, ASP2, ASP3 and DPAP1. PKG is conserved between *B. divergens*, *P. falciparum* and *T. gondii* and has a well-characterized role in egress and invasion in these latter species (Diaz et al., 2006; Donald et al., 2006). PKAc1 suppresses egress in *T. gondii* but is instead required for invasion by *P. falciparum*. The CDPK family of proteins are also known to be required for *P. falciparum* and *T. gondii* egress and invasion, although the direct orthologs do not necessarily have the same function between species. PLPs are required for host cell permeabilization and egress in *T. gondii*, *Plasmodium* spp. sexual stages and *B. bovis* (Deligianni et al., 2018; Deligianni et al., 2013; Kafsack et al., 2009; Paoletta et al., 2021; Wirth et al., 2014). PfDPAP family members, PfPMX, PfPMIX and TgASP3 are required for egress and/or invasion in *P. falciparum* or *T. gondii* (Dogga et al., 2017; Ghosh et al., 2018; Lehmann et al., 2018; Nasamu et al., 2017; Pino et al., 2017; Suárez-Cortés et al., 2016).

To develop a stable transfection system for *B. divergens*, multiple transfection methods and selection drugs with established resistance markers were tested for stable transfection of a GFP reporter plasmid (Figure S3A and C) (summarized in Table S3). All further transfections used Amaxa nucleofection of isolated merozoites and Blasticidin-S selection. Next, a CRISPR/Cas9 system was generated to introduce an HA tag, as well as the *glmS* riboswitch and a destabilization domain (DD) inducible knockdown systems, in series to the 3’ end of each gene (Figures 3A and S3B) (Armstrong and Goldberg, 2007; Prommana et al., 2013). Parasites reached >1% parasitemia 12-16 days after transfection with >90% editing for all constructs, except PKAc1, PLP1 and PLP3 which we were unable to tag (Figure S3G, data not shown). PKG-HA-DD-glmS parasites were initially used to determine the effectiveness of the knockdown systems. Induction of the DD system, which induces protein degradation, generated a more rapid and stronger reduction of protein levels that was observable by 6 hrs (Figure 3B). The glmS system, which destabilizes mRNA, resulted in slower knockdown that was first observable by 24 hrs, however, achieved a strong knockdown by 48 hrs (Figure 3B). Combining both systems resulted in a stronger knockdown than either alone (Figure 3B). This pattern was reflected in the effect of these knockdown systems on parasite growth when targeting PKG, CDPK4, ASP2 or ASP3, with the double knockdown producing the strongest defect in all lines, whereas the DD or glmS were not sufficient in all genes (Figure 3C). DPAP1, PKAc2, CDPK5 and CDPK7 knockdown did not affect proliferation (Figure 3C, data not shown).

**Figure 3.**
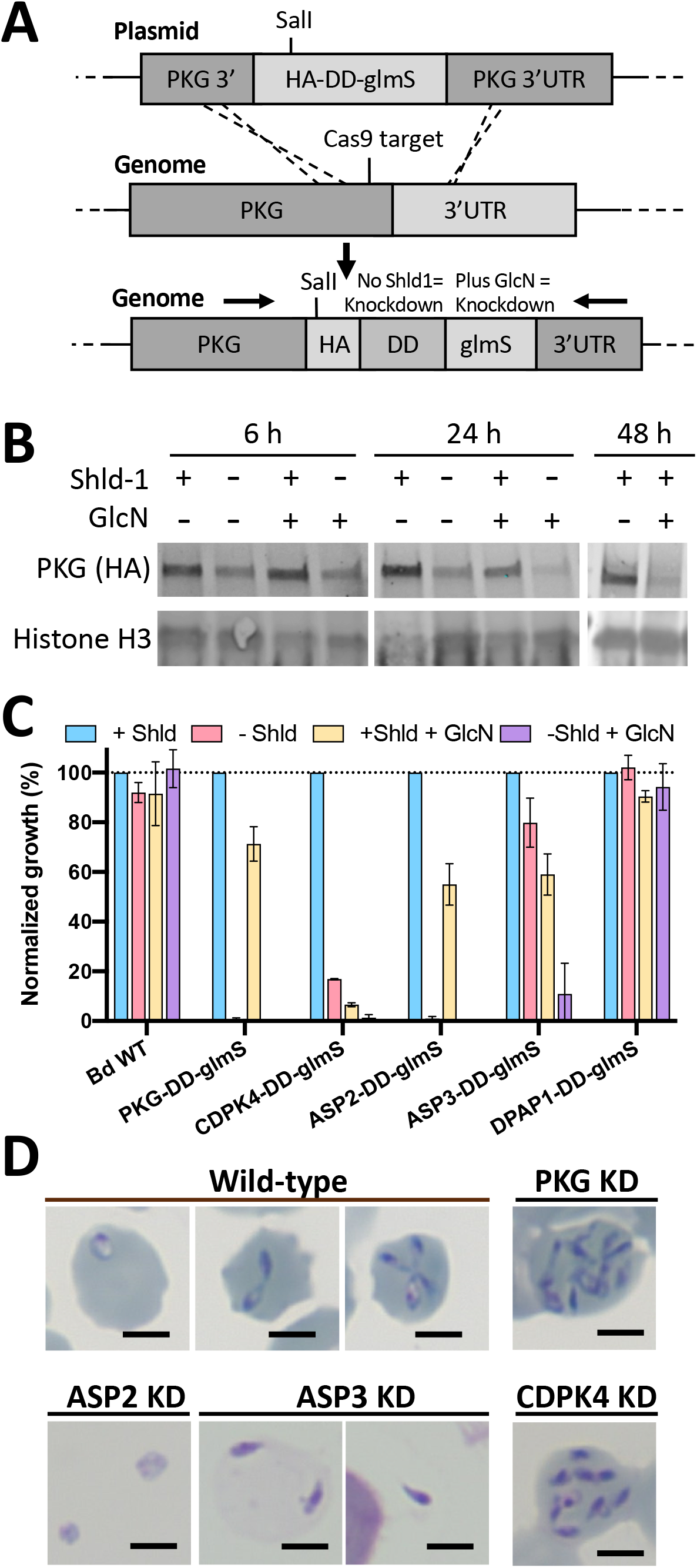
Inducible knockdown reveals an essential role for PKG, CDPK4, ASP2 and ASP3. **(A)** CRISPR/Cas9 is used to introduce a HA-DD-glmS or HA-glmS tag to the 3’ end of *pkg* by homologous recombination. The same approach was used for all genes **(B)** Western blot analysis showing DD and glmS induced knockdown of PKG over time. **(C)** Growth of knockdown parasites over 3 days. Data are the mean ± SD of three biological experiments performed in technical triplicate. **(D)** Giemsa stains showing phenotypes after 24-48 h of knockdown (KD). The scale bar is 3 µM.

### PKG and CDPK4 are critical for egress and their knockdown results in continued intracellular replication

We performed knockdown of PKG and CDPK4 following synchronized invasion to understand when the block in proliferation occurs. PKG or CDPK4 knockdown parasites continued to replicate within a single iRBC, to form up to 16 parasites per RBC by 40 hpi (Figures 3D and 4A). The fraction of parasites that replicate to >4 per iRBC is dependent on the degree of knockdown (Figure 5A). Knockdown of PKG prevented induced egress by 8-Br-cGMP, further supporting that 8-Br-cGMP induces egress through activation of PKG (Figure S3D). The reduced susceptibility to 8-Br-cGMP mediated induced egress in PKG-HA-DD-glmS parasites without inducing knockdown compared to WT parasites is likely due to a partial knockdown in the absence of induction (Figure S3D).

**Figure 4.**
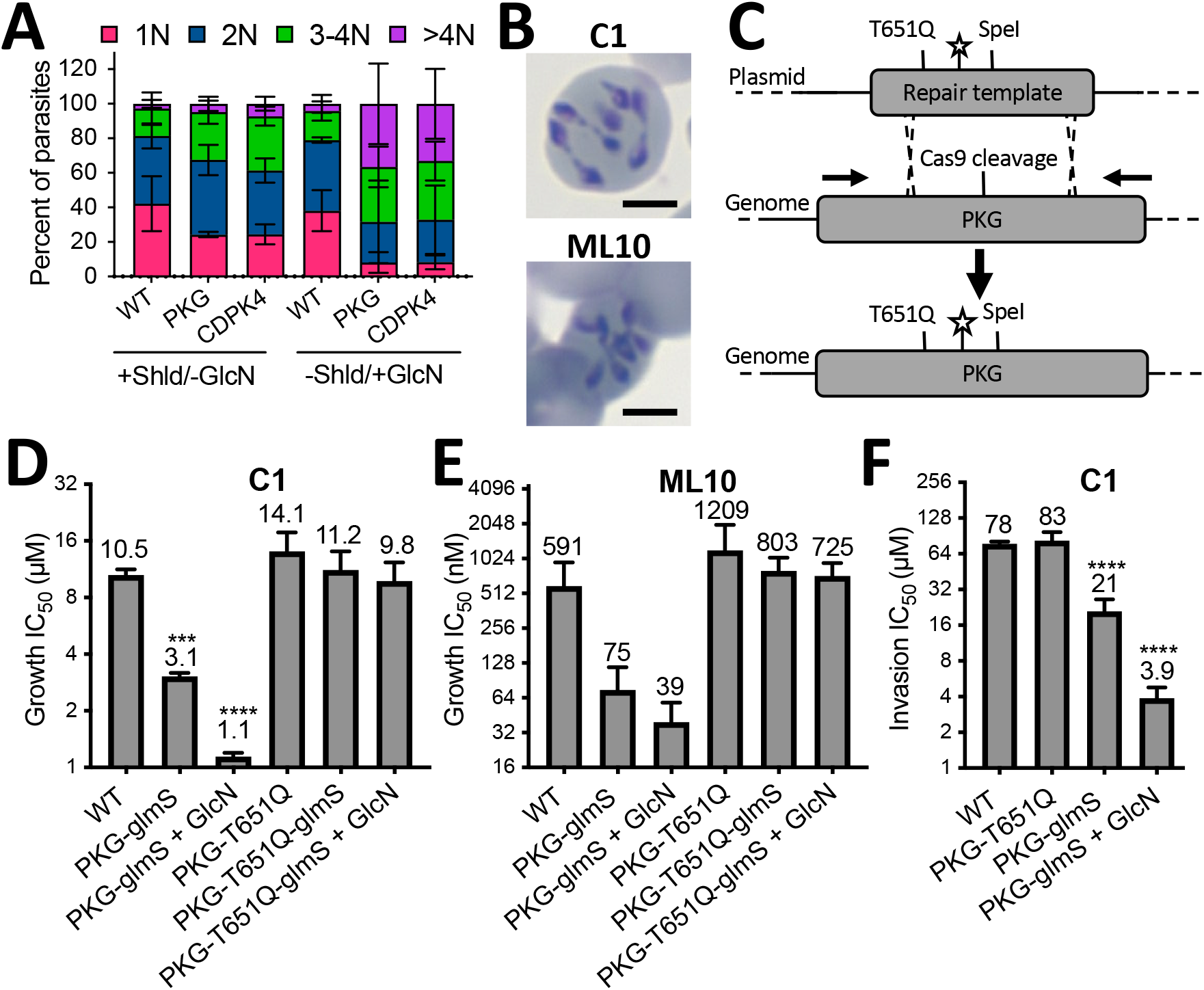
PKG and CDPK4 are required for egress and invasion, and PKG is a druggable target. **(A)** The number of parasites per iRBC assessed by flow cytometry after knockdown (− Shld/+GlcN) of PKG or CDPK4 for 48 h. **(B)** Wild-type parasites treated with 20 µM C1 or 5 µM ML10 for 48 hrs. **(C)** CRISPR/Cas9 is used to introduce a putative drug resistance mutation into PKG (T651Q), a silent shield mutation (star) and a silent SpeI restriction site. **(D-E)** IC_50_ of Compound 1 and ML10 for proliferation **(D-E)** or invasion **(F)**. For (A,D-F), data is the mean ± SD of three biological experiments performed in technical triplicate. The scale bar is 3 µM.

**Figure 5.**
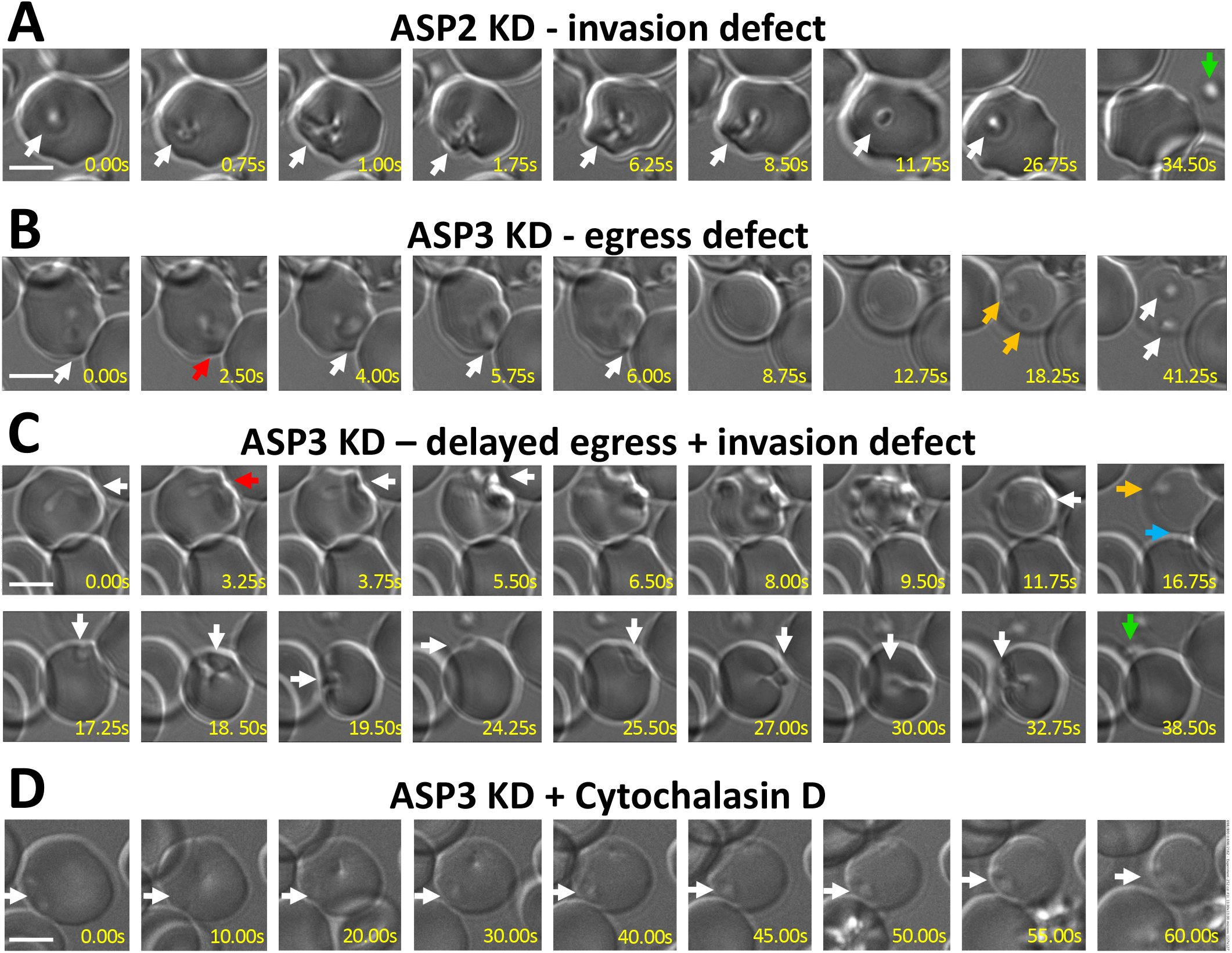
ASP2 and ASP3 are required for egress and/or invasion. **(A-D)** Time-lapse microscopy of 8-Br-cGMP induced egress with knockdown of ASP2 **(A)**, ASP3 **(B,C)** or ASP3 combined with cytochalasin D-treatment **(D)**. Arrows indicate as follows: White – follows the parasites. Red – initial RBC deformation by the intracellular parasite. Orange – failed egress from a permeabilized cell. Blue – successful egress. Green – failed invasion and parasite detachment. The scale bar is 5 µM.

### PKG is an essential and druggable target required for egress and invasion

Significant effort has been invested to develop specific inhibitors of apicomplexan PKG and CPDKs (reviewed in (Arendse et al., 2021; Baker et al., 2020; Hui et al., 2015; Van Voorhis et al., 2017)). To demonstrate that PKG is a druggable target in *B. divergens* we used the apicomplexan PKG inhibitor compound 1 (C1), and ML10 a potent and highly specific *P. falciparum* PKG inhibitor with *in vivo* activity (Baker et al., 2017; Gurnett et al., 2002). Treatment with C1 or ML10 resulted in the same continued intracellular replication as was observed with PKG knockdown (Figure 4B). In contrast to *B. divergens*, *B. bovis* almost exclusively divide once per lytic cycle to form two parasites before egress. However, treatment with C1 caused some parasites to replicate twice to form 4 daughter parasites before egress (Figure S3E), although at a lower rate than *B. divergens,* which was not observed previously (Lau et al., 2013). To determine the specificity of C1 and ML10 for BdPKG, CRISPR/Cas9 was used to introduce a putative resistance mutation (T651Q) equivalent to the gatekeeper mutations that confer resistance in *P. falciparum* and *T. gondii* (Figure 4C)(Donald et al., 2006; McRobert et al., 2008). A 500 bp region within a plasmid was used as a repair template, achieving ∼50% editing (Figure S3G). Two of four edited clones from parasites transfected with the 500 bp repair template were not edited at the more distal positions from the Cas9 cleavage site (24 vs 5 bp) (positions 1-3 of Figure S3F). In 6 of 7 transfections tested, parasites reverted to blasticidin-S sensitive after drug pressure was removed, suggestive of the CRISPR/Cas9 plasmid being lost. We utilized the ability to perform sequential transfection with the same resistance marker to generate double mutants, containing the T651Q mutation and the glmS knockdown system (PKG-T651Q-glmS). A 3- and 10-fold for C1, and 8- and 15-fold for ML10, reduction in the IC_50_ was observed between WT compared to uninduced or induced PKG-glmS parasites, respectively (Figures 4D-F). The difference between WT and uninduced PKG-glmS parasites is likely due to partial knockdown even in the absence of induction. A trending, but not statistically different, shift in the IC_50_ was observed between WT and the PKG-T651Q parasites (10.5 vs 14.1 µM for C1, and 591 nM vs 1209 nM for ML10), whereas the same mutation in knockdown parasites restores susceptibility to near WT levels (Figure 4D-F). Together these results support a model in which C1 and ML10 are able to target PKG, however, at concentrations required for killing *B. divergens* there are secondary targets contributing to growth inhibition. CDPK4 is the only other *B. divergens* CDPK that shares the same small gatekeeper residue as PKG and is the most likely secondary target in the parasite (Figure S3I). The ability to rapidly and specifically inhibit PKG with C1 when combined with partial knockdown was used to investigate the role of PKG during RBC invasion. Knockdown was induced with the glmS system for 48 hrs prior to isolation of free merozoites. PKG knockdown increased the sensitivity to C1 invasion inhibition by 20-fold, demonstrating a requirement for PKG activity during host cell invasion (Figure 4F). Together these data demonstrate that PKG is an essential and druggable protein in *B. divergens* that is required for both egress and invasion.

### ASP2 and ASP3 are required for egress and invasion

ASP2 and ASP3 knockdown resulted in an increased number of free merozoites in culture (Figure 3D). In the ASP3 knockdown culture, clusters of parasites were also observed in lightly stained RBCs (Figure 3D, ASP3 left panel). To further define these phenotypes, induced egress in knockdown parasites was observed by video microscopy. ASP2 knockdown parasites displayed no obvious defect in egress and were able to bind to and strongly deform the host RBC, however, were unable to complete invasion and eventually detached from the cell (Figure 5A, Video 6). Egressing ASP3 knockdown parasites significantly deformed the host cells from within for ∼7 seconds (compared to <0.5 s in WT parasites) before the eventual permeabilization of the host cell, suggesting a defect in release or activity of secreted lytic factors required for egress. Once the host cell was permeabilized, the parasites were not always motile, remaining trapped in the permeabilized cell (Figure 5B). The parasites that were motile and escaped the host cell were able to bind to and deform the RBC, however, unlike ASP2 knockdown parasites that typically bind strongly to a single host cell, ASP3 parasites maintained gliding motility over the surface of the cell, often contacting multiple host cells, but were rarely able to complete invasion (Figure 5C, Video 7). These results suggested that ASP3 knockdown parasites may be utilizing motility to disrupt the host cell in the absence of lytic factors, as has been observed with TgPLP1 knockout parasites (Kafsack et al., 2009). To test this hypothesis, ASP3 knockdown parasites were treated with cytochalasin D to inhibit gliding motility and induced to egress. These parasites were still able to lyse the host cell albeit over a significantly longer time period (∼60 s), suggesting that parasites were secreting active lytic factors at a reduced rate (Figure 5D, Video 8). In *P. falciparum*, incomplete inhibition of PfPMX results in parasites lysing the PVM, whereas complete inhibition blocked PVM rupture (Favuzza et al., 2020; Pino et al., 2017). Further work will be required to determine if complete inhibition of ASP3 in *B. divergens* would block RBC lysis. Together, these results demonstrate that ASP2 is required for normal invasion at a step later than that of ASP3, consistent with a rhoptry function. ASP3 is required for efficient host cell permeabilization, escape from the permeabilized cell and invasion into the next host cell consistent with a microneme function.

## Discussion

The majority of molecular research in apicomplexan parasites has been limited to *P. falciparum, P. bergheii* and *T. gondii*. Transient or stable transfection systems exist for *Babesia* spp., *Theileria* spp., *Cryptosporidum* spp., *Eimeria* spp., *Sarcocystis* spp. and *Neospora* spp. (reviewed in Suarez et al. (2017)). However, the number of studies and the range of available tools and resources remains comparatively limited in these organisms. Additional studies of a wider range of apicomplexan parasites will expand our knowledge of conserved and unique biological mechanisms of apicomplexan parasitism. Here, we establish *B. divergens* as a genetically tractable *in vitro* model to study *Babesia* spp. cell biology. The synchronous *B. divergens* bulk transcriptome and single-cell transcriptome will serve as a resource for the study of *Babesia* spp. biology as well as for further comparative studies of apicomplexan biology.

### The *B. divergens* egress model

As egress is a unique aspect of apicomplexan parasitism compared to their hosts, they offer novel druggable targets. The overall process of egress is similar between *Plasmodium* spp. and *T. gondii*, however, there are notable differences in their cell biology (e.g. differences in division mechanisms) and in the host cell niche (*T. gondii* resides in nucleated cells, while the asexual stages of *Plasmodium* spp. reside in enucleated RBCs), that place unique pressures on the parasite. While the signaling and molecular mechanisms differ in some aspects, we have found that egress of *B. divergens* at the cellular level closely resembles that of *T. gondii* and has several notable differences to *Plasmodium* spp. asexual stage egress despite being evolutionarily closer to *Plasmodium* spp. and sharing the same RBC niche. Here, we propose a model of *B. divergens* egress in which the parasite responds to an initial signal (intrinsic or extrinsic) to induce the PKG- and CDPK4-dependent egress signaling pathway (Figure 6). The egress signaling pathway leads to the release of lytic factors (e.g. PLPs (Paoletta et al., 2021)) into the host cell, which damages the RBC membrane but leaves the cytoskeleton relatively intact. *B. divergens* parasites then utilize gliding motility to escape the permeabilized host RBC and seek a RBC to invade.

**Figure 6.**
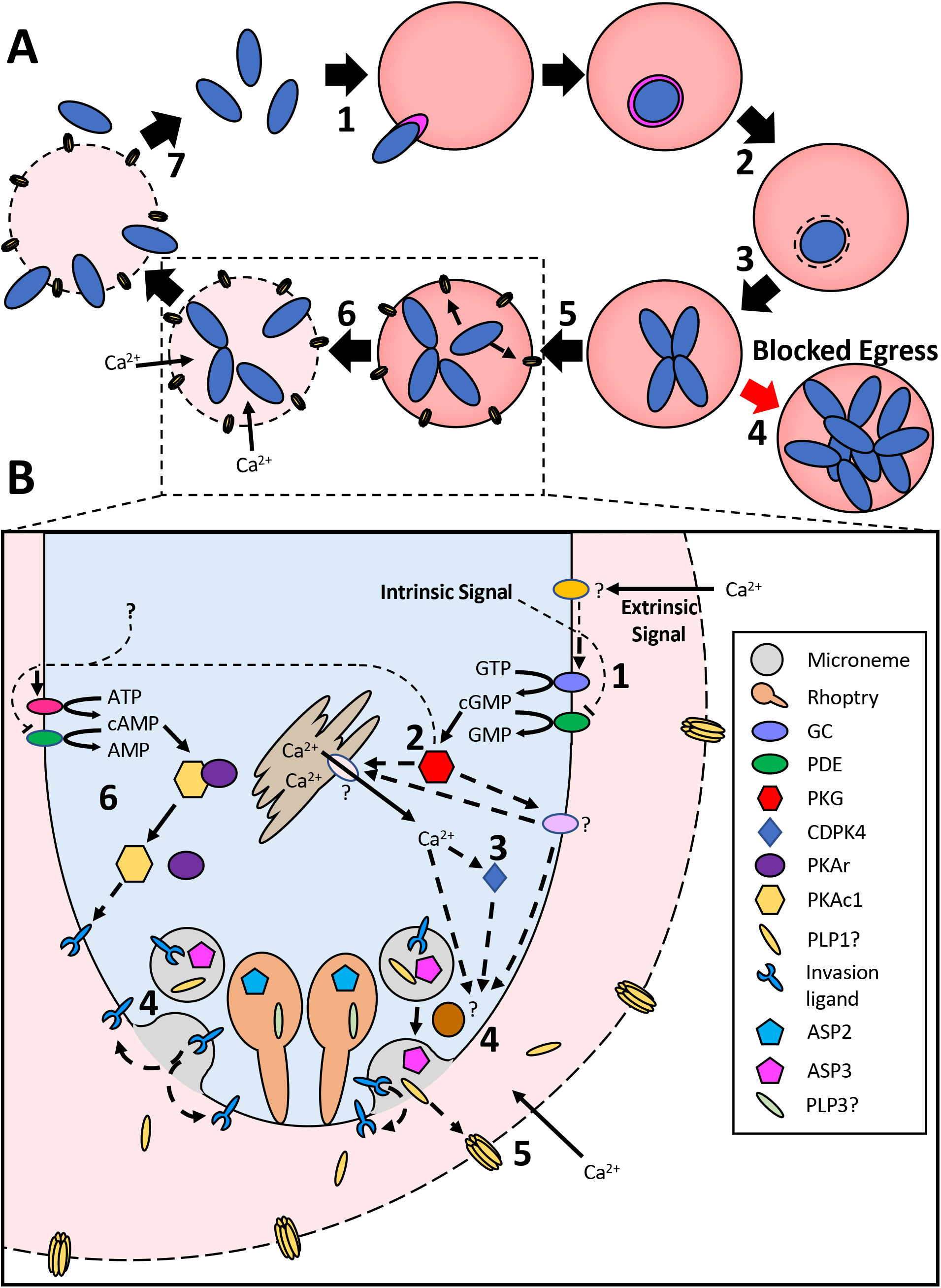
Model of the Babesia divergens lytic cycle. **(A)** 1. *B. divergens* merozoites invade a RBC and are surrounded by the newly formed PVM. 2. The PVM is degraded. 3. One or two replication cycles occur to produce 2-4 parasites. 4. In the absence of an egress signal, the parasite continues to divide within a single RBC. 5. An unidentified intrinsic signal triggers egress in one or more parasites. 6. Secreted parasite lytic factors permeabilize the RBC membrane leading to an influx of calcium. 7. Calcium acts as an extrinsic signal to trigger egress and gliding motility in the remaining parasites to escape the permeabilized RBC. **(B)** 1. An extrinsic signal (calcium) or an intrinsic signal activates guanylate cyclase (GC) and/or inhibits a phosphodiesterase (PDE). 2. The increased cGMP levels activate cGMP-dependent kinase (PKG), which leads to the release of calcium and putatively activates PKA and the glideosome. 3. Calcium, CDPK activity and potentially lipid signals lead to the fusion of micronemes to that parasite plasma membrane. 4. The contents of the micronemes are released into the RBC cytosol, including lytic factors (putatively PLP1), and invasion ligands onto the parasite surface 5. Lytic factors act to permeabilize the RBC membrane. The permeabilized RBC membrane allows an influx of calcium, acting as a positive feedback loop. 6. Unknown signals, potentially including PKG, lead to the activation of adenylate cyclase (AC) and/or inhibition of a PDE, which increases cAMP levels. The increased cAMP activates PKAc1, which is required for invasion, putatively through the phosphorylation of invasion ligands including AMA1. Solid lines represent pathways with a high confidence based on available data for *B. divergens*. Dashed lines and proteins marked with a “?” indicate processes based primarily on the RNAseq data from this paper and orthology to *P. falciparum* or *T. gondii*.

### Molecular mechanism: similarities and deviances

#### Intrinsic and extrinsic factors that induce egress

*T. gondii* is able to sense phosphatidic acid (PA) and low pH, which act as intrinsic signals of parasite density to induce egress (Bisio et al., 2019; Bullen et al., 2016), as well as extracellular potassium, calcium and serum albumin, which act as extrinsic signals of host cell damage to induce egress (Brown et al., 2017; Brown et al., 2016; Gaji et al., 2011; Moudy et al., 2001; Vella et al., 2021). *P. falciparum* is also able to sense lipids, including phosphatidylcholine (PC) and diacylglycerol (DAG), as well as potassium to enhance egress and microneme secretion, although their roles in the parasite are less clearly defined (Paul et al., 2020; Singh et al., 2010). Since *Plasmodium* spp. egress is strictly required at the end of each replication cycle, the initial egress signal is likely to be linked to the cell cycle, such as expression of the PKG signaling pathway, however, this remains to be demonstrated. The only egress signal identified in *B. divergens* is extracellular calcium, but not potassium or sodium, which acts as an extrinsic signal of host cell damage (Figure 1F, Video S4). The extracellular calcium-sensing pathway likely signals through PKG, since the addition of 8-Br-cGMP can bypass it to induce motility (Figure 1F, Video S2).

#### The egress signaling pathway

In *T. gondii* and *P. falciparum*, these initial signals converge to either activate GC or inhibit PDE and thus raise cGMP levels, which in turn activates PKG. In *T. gondii*, PKAc1 suppresses cGMP levels to prevent premature egress, putatively by activating a PDE, whereas in *P. falciparum* PKAc1 is required for invasion (Dominicus et al., 2021; Flueck et al., 2019; Jia et al., 2017; Moss et al., 2021; Patel et al., 2019; Wilde et al., 2019). Inhibition of PKA using H89 blocked *B. divergens* invasion, as well as induced low levels of egress. Similarly, treatment with 8-Br-cAMP inhibited induced egress, suggesting that *B. divergens* PKA is required for both egress and invasion (Figure 1G, S1D and S1G). The two PKAc orthologs in *B. divergens* could be involved in separate roles during egress and invasion, however, only PKAc1 displayed stage-specific expression. We were unable to knockdown PKAc1 and validation of its function will require further study. *P. falciparum* uses protein phosphatase 1 (PP1) to increase cGMP levels, putatively through dephosphorylation of the large LT-GC protein (Paul et al., 2020). *B. divergens* has a PP1 ortholog which is expressed in blood stages but displays no stage-specific expression and remains uncharacterized. In *P. falciparum* and *T. gondii*, PKG activates PI-PLC, which generates lipid second messengers. These eventually lead to the release of calcium from intracellular stores that are required for microneme/exoneme secretion. We have demonstrated that PKG is a central component of the *B. divergens* egress signaling pathway that is both necessary and sufficient to induce egress (Figures 1D-F and S3D). Small molecules targeting the lipid signaling pathway, including propranolol and U73122, had little to no effect on *B. divergens* egress, while R59022 enhanced egress rather than inhibited it like it does in *T. gondii* and *P. falciparum*. To determine if these differences are due to technical reasons or divergence of the signaling pathways, further genetic and chemical studies in *Babesia* spp. Will be needed. Downstream of PKG and calcium release, CDPKs are required in *P. falciparum* and *T. gondii* egress. *B. divergens* only expresses three CDPKs in *in vitro* culture, CDPK5, CDPK7 and CDPK4, of which only CDPK4 displayed stage-specific expression indicative of a role in egress (Figure 2E and 2G, Table S1). Knockdown of CDPK4 or chelation of calcium with BAPTA-AM, inhibited 8-Br-cGMP induced egress, however, release of calcium with the ionophore A23187 did not induce egress or motility in *B. divergens*. Together these results support that calcium release and CDPK4 activity are required, but not sufficient for egress, suggesting they likely act downstream of PKG in the same pathway, although we note that a previous study found that A23187 could induce egress in *B. bovis* (Mossaad et al., 2015).

The ‘decision’ of when to egress appears to occur proximal to the point of activation of the PKG/CDPK4 signaling pathway. Inhibition of PKG and CDPK4 as well as chelation of extracellular calcium all result in continued intracellular replication in *B. divergens* and/or *B. bovis* (Figures 3D, 4A and S3E) (Cursino-Santos et al., 2017; Pedroni et al., 2016). Inhibition of either the signaling pathway or the downstream lytic factors produces a similar phenotype in *T. gondii* (Donald et al., 2006; Kafsack et al., 2009). Indeed, PLP1 gene deletion in *B. bovis* resulted in iRBCs containing >2 merozoites (Paoletta et al., 2021). This implies that *T. gondii* and *Babesia* spp. lack a checkpoint to exit the cell cycle and that the default pathway is to continue replication. Inhibition of CDPK5 blocks *P. falciparum* egress and replication, however, the transcriptional profile of stalled parasites continues to progress, suggesting that they may also lack a checkpoint at the transcriptional level (Dvorin et al., 2010).

### Cell lysis and motility

*T. gondii* uses PLP1 and LCAT to permeabilize the PVM and host cell membrane (Kafsack et al., 2009; Schultz and Carruthers, 2018), whereas the *P. berghei* orthologs (PLPs and PbPL) are not essential for asexual stage egress and no other mechanisms have yet been identified (Bhanot et al., 2005; Sassmannshausen et al., 2020). We identified *B. divergens* PLP1 (ortholog of TgPLP1) as a possible lytic factor that displays an expression profile matching that of microneme proteins (Figure 2E and G). No orthologs of the phospholipase TgLCAT/PbPL were identified in the *B. divergens* genome. The first step in egress in *Babesia* spp. has been re-wired to degrade the PVM immediately after invasion. We identified PLP3 (no direct ortholog) as having an expression profile consistent with rhoptry localization and could be secreted into the newly formed vacuole to mediate its degradation, however, this will require further genetic studies (Figure 2 E and G). *P. falciparum* SERA6 is required for disruption of the RBC cytoskeleton and is the putative target of E64 (Thomas et al., 2018). *Babesia* spp. do not have an ortholog of any SERA proteases and do not degrade the RBC cytoskeleton, which is consistent with the limited inhibition of egress by cysteine protease inhibitors.

Many of the proteins involved in egress and invasion are known to be proteolytically matured in *T. gondii* and *Plasmodium* spp. In *P. falciparum*, proteolytic maturation in the rhoptries and micronemes is done in part by PfPMIX and PMX, respectively. In contrast, *T. gondii* only contains one ortholog, TgASP3, which is localized in a post-Golgi compartment and is able to mature a subset of microneme and rhoptry proteins. It is likely that BdASP2 and BdASP3 also act to mature a range of proteins required for egress and invasion. Their putative localization based on transcriptomics and their knockdown phenotypes suggest that BdASP2 and BdASP3 are functionally orthologous to PfPMIX and PfPMX, respectively. Interestingly, the invasion phenotype differs between BdASP2 and BdASP3, with ASP2 knockdown parasites binding to a single host cell where they remain attached for an extended period, suggesting they may reorientate, but are not able to undergo the final step of invasion which requires rhoptry proteins (e.g. RON2 (Lamarque et al., 2011; Treeck et al., 2009; Yap et al., 2014). In contrast, BdASP3 knockdown parasites that are able to egress, bind to, and strongly deform, the RBC but remain motile for an extended period of time and often interact with multiple RBCs before losing motility. This phenotype suggests that ASP3 is required for maturation of proteins that are required for reorientation and/or anchoring of the apical end of the parasite to the RBC prior to rhoptry release (Weiss et al., 2015a). The efficiency of egress in BdASP3 knockdown parasites was reduced, with strong deformation of the RBC for several seconds prior to RBC permeabilization in contrast to the rapid process in wild-type parasites. Despite this strong physical deformation of the RBC prior to lysis, it is not necessary for the lysis as demonstrated through treatment with cytochalasin D (Figure 5D), suggesting the secreted lytic factors are still present but have reduced activity or levels.

### Egress and invasion contain multiple druggable targets

Small molecules have been developed that target the kinases and proteases required for egress of *Plasmodium* spp. and *T. gondii*. Identification of shared parasite targets of these compounds in *Babesia* spp. and other apicomplexan parasites will help leverage drug development efforts for malaria that have significantly more resources. Here, we have developed a transfection and CRISPR/Cas9 and inducible knockdown systems to modify the *B. divergens* genome. Through knockdown and the introduction of resistance mutations, we identified C1 and ML10 as inhibitors of PKG, however, at concentrations required for killing it is likely that other parasite molecules are also inhibited (Figure 5D-E). Using chemical and genetic methods in combination to target a protein can overcome the slow nature of inducible knockdown systems (∼6 hrs for PKG-DD-glmS, Figure 3B) and potential lack of specificity of chemical inhibition to validate specific targets. This was used to demonstrate that PKG is required for invasion (Figure 4F). We have also generated inducible knockdown parasites for CDPK4, ASP2 and ASP3, the orthologs of which are all areas of active drug development (reviewed in (Caldas and De Souza, 2018; Singh and Chitnis, 2017). The same methods could be applied to these, and other, proteins to aid in future drug development for *Babesia* spp.

Here, for the first time we have established a molecular framework for the events surrounding *Babesia* spp. egress. Nevertheless, many questions remain unanswered. While we have identified a set of genes that are putatively involved in egress or invasion, we do not know many of the molecular mediators of egress, how *Babesia* spp. selectively lyse the PVM, the role of lipid signaling or the source of intracellular calcium utilized during egress. The initial natural signal to induce egress and how *Babesia* spp. regulate egress to occur after one or more replication cycles have also not been identified. With the cellular and genetic tools developed here, future studies will be able to answer these questions and reveal unique biology of *Babesia* spp. as well as conserved processes throughout apicomplexa.

## RESOURCE AVAILABILITY

### Lead contact

Further information and requests for resources and reagents should be directed to and will be fulfilled by the lead contact, Manoj Duraisingh.

### Materials availability

All unique reagents in this study are available from the lead contact. Any additional information required to reanalyze the data reported in this paper is available from the lead contact upon request.

### Data and code availability

Single-cell RNAseq and bulk RNAseq data have been deposited at NCBI Sequence Read Archive (SRA) and are publicly available as of the date of publication. Accession numbers are listed in the key resources table. Microscopy data reported in this paper will be shared by the lead contact upon request. All code has been deposited at GitHub. DOIs are listed in the key resources table.

## EXPERIMENTAL MODEL AND SUBJECT DETAILS

### Parasite strains

*Babesia divergens* strain Rouen 1987, kindly provided by Kirk Deitsch and Laura Kirkman (Weill Cornell Medical College), was cloned by limiting dilution (BdC9) and was maintained in purified Caucasian male O+ human RBCs (Research Blood components). The *Babesia bovis* strain MO7 was kindly provided by David Allred of the University of Florida and maintained in purified bovine RBCs (hemostat). Parasites were cultured in RPMI-1640 media supplemented with 25 mM HEPES, 11.50 mg/l hypoxanthine, 2.42 mM sodium bicarbonate, and 4.31 mg/ml AlbuMAX II (Invitrogen), at 37° C in a 1% oxygen, 5% carbon dioxide and 94% nitrogen environment.

## METHOD DETAILS

### Induced egress

To measure egress, mixed stage parasites at 2% final HCT, 10-15% parasitemia in a total volume of 40 µL in the presence of the stated drug, were incubated at 37° C for one hour in a 96 well plate for 1 hr. Where not otherwise stated, 8-Br-cGMP was used at a final concentration of 500 µM. For inhibition of induced egress, parasites were first incubated with the stated compound at 1.33x concentration for 15 min at 37° C, prior to induced egress with the addition of 8-Br-cGMP to a final concentration of 500 µM and 1x inhibitor concentration. A final concentration of 100 µg/ml of heparin was added to prevent reinvasion. After 1 hr, the parasites were washed 3x with PBS and stained with 1:5000 Syber green II. Parasitemia was determined by flow-cytometry (Macs quant, Miltenyi) and analyzed in FloJo. Induced egress was calculated as the drop in parasitemia of the treated parasites as a percentage of the RPMI only sample.

### Live microscopy

All videos were taken on a Zeiss Axio Observer using a 60x oil immersion lens inside a chamber heated to 37° C. Prior to imaging, parasites were allowed to settle on the bottom of a glass bottom slide (Ibidi, Cat#80827/81817) at 37° C in a 5% CO_2_ incubator for 10-15 min. Small molecule inhibitors were included in this incubation as necessary. Immediately prior to imaging, the media was removed and replaced with the RPMI/IC/EC with 500 µM 8-Br-cGMP, and any small molecules inhibitors being tested. For IC/EC experiments, the final buffer contained 0.0075% (w/v) saponin (Calbiochem, Cat#558255) to lyse the RBC. Alexa Fluor^TM^ 488 Phalloidin (Invitrogen, Cat#A12379) was added to RPMI at a concentration of 1/150 where stated. All images were taken within 20 minutes of re, maged as per the *B. divergens* protocol. Images were processed in Zen 2 (Zeiss) and ImageJ/Fiji.

### Isolation of free merozoites for invasion assays, synchronization and transfection

∼1-2 ml of packed iRBCs at 20-30% parasitemia was used to isolate free merozoites using a modified protocol from (Cursino-Santos et al., 2016). Briefly, the iRBC was resuspended to 10% HCT in RPMI and passed through two 1.2 µM filters. The isolated merozoites and RBC debris were pelleted at 3000 x g for 3 min and the supernatant was removed. For invasion assays, the merozoites were resuspended in RPMI and added to a 96 well u-bottom plate containing 1.33x final drug concentration (30 µl total) and incubated at 37° C for 10 min. Fresh RBCs were then added to a final of 2% HCT and 1x drug concentration (40 µl total). The plate was then incubated at 37° C shaking at 600 rpm for 20 min. The assay was stopped by washing the parasites three times with 200 µL PBS, 400 x g for 2 min or by adding 200 µl 4% paraformaldehyde. Parasitemia was determined by flow cytometry as per induced egress section above. For synchronization, isolated merozoites were resuspended to a final volume of 1 ml of 20% HCT RBCs in RPMI and allowed to invade for 20 min shaking at 600 rpm and 37° C. Parasites were then washed 3x with 10 ml RPMI at 400 x g to remove free merozoites and cell debris. 100 µg/ml heparin was added to prevent re-invasion throughout the time course when stated in the figure legend.

### Plasmid construction

The sequences of all primers and synthesis products used in this study are found in table 4. The bi-directional promoter between the EF1alpha (Bdiv_030590) and LON peptidase (Bdiv_030580c) was amplified using primers BE-8 and BE-9, and cloned into XhoI/BamHI sites of pPfEF-GFP-BSD (EF1alpha side drives GFP/Cas9 expression). The DHFR (Bdiv_030660, primers BE-21/22) and HSP90 (Bdiv_037120c, primers BE-32/33) 3’UTRs were cloned into EcoRI/HindIII and SpeI/NotI sites, respectively, for the selection marker or GFP/Cas9, respectively. Cas9 was amplified from pDC2-Cas9 using primers BE-125/125 and cloned into the XhoI/SpeI sites in place of GFP. The U6 promoter, bbs1 sites (for the guide), guide tracer/scaffold, U6 terminator and PKG-T651Q repair template were synthesized by IDT (Synthesis 1) and cloned into the EcoRI site by gibson assembly to make the pEF-Cas9-PKG-T651Q plasmid. For PKG, guide oligos were phosphorylated, annealed and cloned into the BbsI sites. For all inducible knockdown lines, the HA-glmS or HA-glmS-DD tags were amplified using primers BE-536/537. The 5’HR (HR1, 3’end of the gene) and 3’HR (HR2, 3’UTR of gene) were amplified using the corresponding HR1 and HR2 primers from Table 4 for each gene. The sgRNA was amplified using the corresponding ‘guide PCR’ primer (e.g. BE-551) and BE-550. The sgRNA, HR1, DD-glmS and HR2 PCR fragments were cloned into the Bbs1/PacI sites of the pEF-Cas9-PKG-T651Q plasmid in a single Gibson reaction.

### Transfection

For transfection of iRBCs using the Biorad Genepulser II, 100 µg DNA in 30 µl combined with 370 ul of cytomix (120 mM KCl, 0.15 M CaCl2, 2mM EGTA, 5mM MgCl2, 10mM K2HPO4/KH2PO4, 25mM HEPES, pH 7.6) and 200 µl iRBCs (∼15% parasitemia). Transfection was carried out with Biorad genepulser II set to 0.31 kV, 950µF. For Amaxa nucleofection of free merozoites, merozoites were isolated from ∼3×10^9^ iRBCs and all supernatant was removed. The pellet, containing free merozoites and cell debris, was resuspended in 100 µl of P3 solution (Lonza) plus 10 µl of water containing 2-10 µg of DNA. For Amaxa nucleofection of intact iRBCs, 10 µl iRBCs at 10% parasitemia was resuspended in 110 µl P3+DNA solution, as per above. Transfection of either free merozoites or iRBCs was carried out in a 4D-Nucleofector System (Lonza), using the FP158 program. After electroporation of free merozoites, the parasite and buffer mixture was immediately transferred to 1 ml of RPMI containing 200 µl packed RBCs and pre-heated to 37° C. Parasites were allowed to invade at 37° C shaking at 600 rpm for 20 min before being washed with 10 ml RPMI to remove the P3 solution and returned to culture. Electroporated intact iRBCs were washed 1x with 10 ml RPMI and returned to culture. Cultures were selected with 15-20 µg of Blasticidin-S.

### Drug sensitivity and proliferation assays

Parasites were diluted to 0.2% parasitemia in 2% hematocrit (100 µl total) in the presence of the stated compounds) in a 96-well U-bottom plate. Parasites were cultured for 72 hours. Parasitemia was measured by flow-cytometry.

### RNAseq analysis

RNA was isolated from parasites using a hybrid protocol of organic extraction combined with column purification. Briefly, parasite pellets were resuspended in 500 µl TRIzol and then extracted with chloroform. The aqueous layer was then purified using Qiagen RNeasy Mini spin-columns following manufacturers protocols. RNA was quantified and normalized into individual wells, and libraries were prepared following the Smart-seq2 protocol (Picelli et al., 2013). Libraries were sequenced on the illumina platform. Bulk synchronous RNAseq analysis workflow: The quality of reads was assessed using FastQC (Version 0.10.1). The reads were trimmed with Cutadapt from Trim Galore package (Version 0.3.7) (http://www.bioinformatics.babraham.ac.uk/projects/trim_galore/). The trimmed reads were mapped against the Bdivergens1802A reference genome (PiroplasmaDB release 46) and assembled with HISAT2 (Version 2.0.5 released)(https://www.ncbi.nlm.nih.gov/pmc/articles/PMC4655817/). SAM files obtained from alignment results were processed using SAMtools (Version 1.4.1) and the relative abundance of transcripts were estimated using featureCounts (https://academic.oup.com/bioinformatics/article/30/7/923/232889). Normalization & Noise removal: Counts per million (cpm) values per gene was calculated using cpm() from edgeR R package (Version 3.24.3) (https://www.bioconductor.org/packages/release/bioc/vignettes/edgeR/inst/doc/edgeRUsersGuide.pdf). Genes with cpm value > 2 in at least 3 samples were maintained for further analysis. Gene counts were normalized and scaled to logarithmic form using edgeR’s TMM method (trimmed mean of *M* values) with DGEList(), calcNormFactors() and cpm() functions. The cpm() parameters were as following: y = DGEList.obj, log=TRUE, prior.count=3, normalized.lib.sizes=TRUE. Batch effected samples were Identified through the analysis of hierarchical clustering and dissimilarities method using the R function hclust() with the default parameters and logCPM expression values. These samples were excluded from further analysis. The bad samples were the ones clustered in single batches and were not similar to the rest of samples based on Euclidean distance metric. PCA Analysis: Principal component analysis (PCA) was run on genes with FC > 1.5 with prcomp() from R. The first 2 principal components (PCs) were chosen for visualizati nalysis: Orthologous genes between *Babesia divergens* 1802A, *Plasmodium falciparum* 3d7, *Toxoplasma gondii* ME49, *Plasmodium berghei* ANKA (all from VEuPathDB release 46) were assembled using blast (version 2.10.0) (https://academic.oup.com/nar/article/36/suppl_2/W5/2505810) and the R package Orthologr (version ‘0.4.0’) with implemented best reciprocal hit method.

### scRNA-Seq work flow

scRNA-Seq data was processed using the 10x cell-ranger pipeline and aligned to the *Babesia divergens* 1802A genome. Counts were subsequently normalized and processed using the R Seurat package. A total of 9450 cells and 3620 genes were retained after removing cells and features with low counts (Seurat parameters: min.cells = 10, min.features = 100, nFeature_RNA > 200 nFeature_RNA < 1200). Dimensionality reduction and clustering analysis were performed using PCA and graph-based KNN as implemented in the Seurat Package. A total of 4 clusters were identified by the KNN algorithm. To make the size of the data manageable, each cluster was downsampled to include 800 cells. Global differential expression performed on each cluster (log(FC) > 1 and adjusted p-value < 0.01) identified 544 differentially expressed genes (173 genes in cluster 0, 81 in cluster 1, 149 in cluster 2, and 141 in cluster 3). Pseudo-time analysis: Pseudo-time analysis was performed on the first two PCA components by fitting a principal curve to the data and orthogonally projecting the cells on the curve. Gene expression curves were then constructed using the pseudo-time and the start of gene expression was identified by cross correlation analysis between the bulk and single cell expression curves. Further details can be found in (Rezvani et al., 2022).

## QUANTIFICATION AND STATISTICAL ANALYSIS

All statistical analysis was performed in Graphpad PRISM 9. IC_50_ values were calculated using the Non-linear regression function (variable slope – four parameters, least squares regression). Statistical significance of IC_50_ changes were determined using ANOVA. All image analysis was performed in Zen 2 (Zeiss) and ImageJ/Fiji. Details of each analysis can be found in the corresponding figure legend.

### KEY RESOURCES TABLE

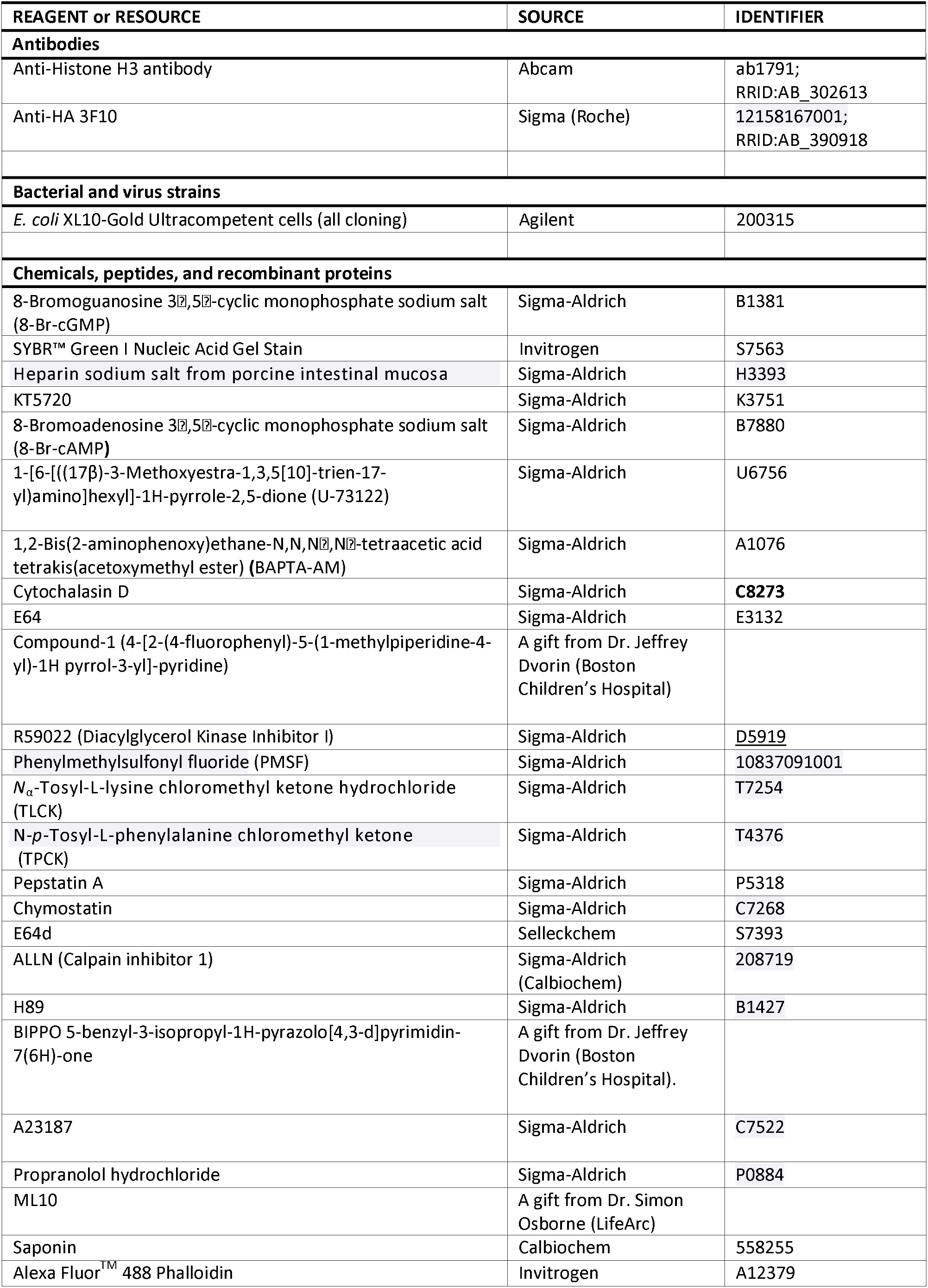
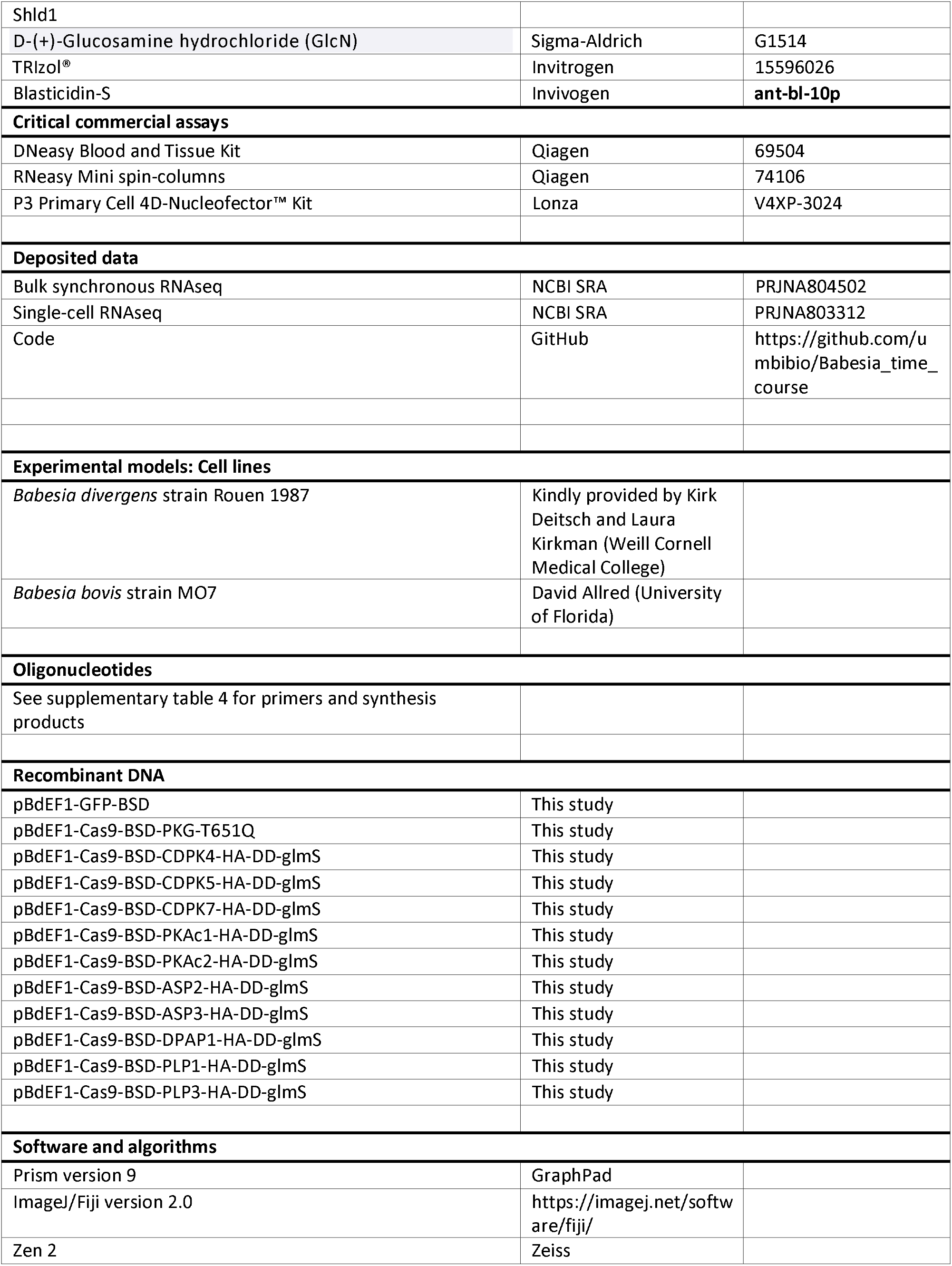
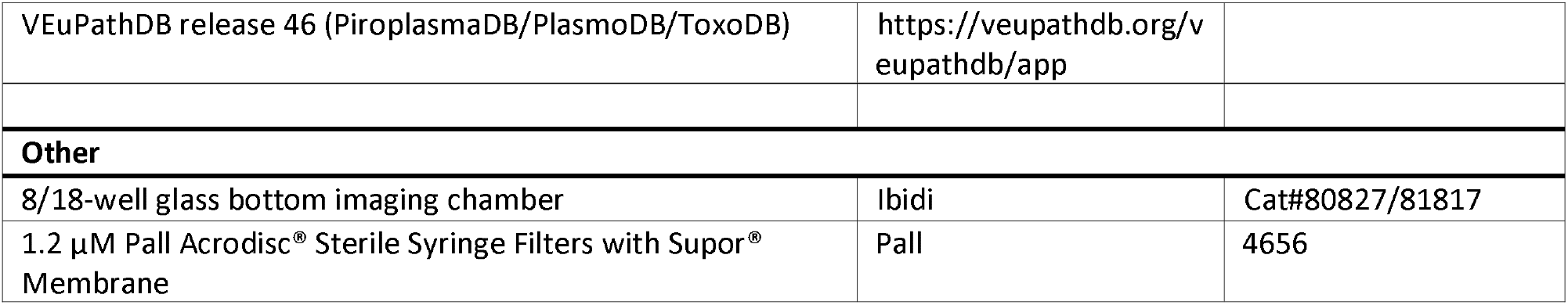

## Supporting information

Video S1. 8-Br-cGMP induced egress

Video S2. EC no calcium plus 8-Br-cGMP

Video S3. 8-Br-cGMP induced egress of Cytochalasin D treated parasites

Video S4. IC and EC with calcium plus saponin

Video S5. IC and EC without calcium plus saponin

Video S6. ASP2 knockdown plus 8-Br-cGMP

Video S7. ASP3 knockdown plus 8-Br-cGMP

Video S8. ASP3 knockdown plus cytochalasin D plus 8-Br-cGMP

Supplementary Table 1. Orthologs of known apicomplexan egress and invasion genes.

Supplementary Table 2. Putative novel egress and invasion genes in B. divergens.

Supplementary Table 3. Summary of selection drugs, resistance markers and transfection methods used for B. divergens.

## Acknowledgment

We would like to thank Dr. Simon A. Osborne (LifeArc) for kindly providing ML10 and Dr. Jeffry D. Dvorin for providing Compound 1 and BIPPO. We would like to thank Dr. Sebastian Lourido for providing A23187 with demonstrated activity in *T. gondii*. This work was supported by grants 1R21AI153945 (MD) and AI150090 (KZ, MJG) from the National Institutes of Health. BE was supported by an American Heart Association (AHA) postdoctoral fellowship (17POST33410556) and a Australian National Health and Medical Research Council postdocotral fellowship (APP1148392). CDK was supported by an AHA predoctoral fellowship (19PRE34380106).

---

**Supplementary Figure 1.**
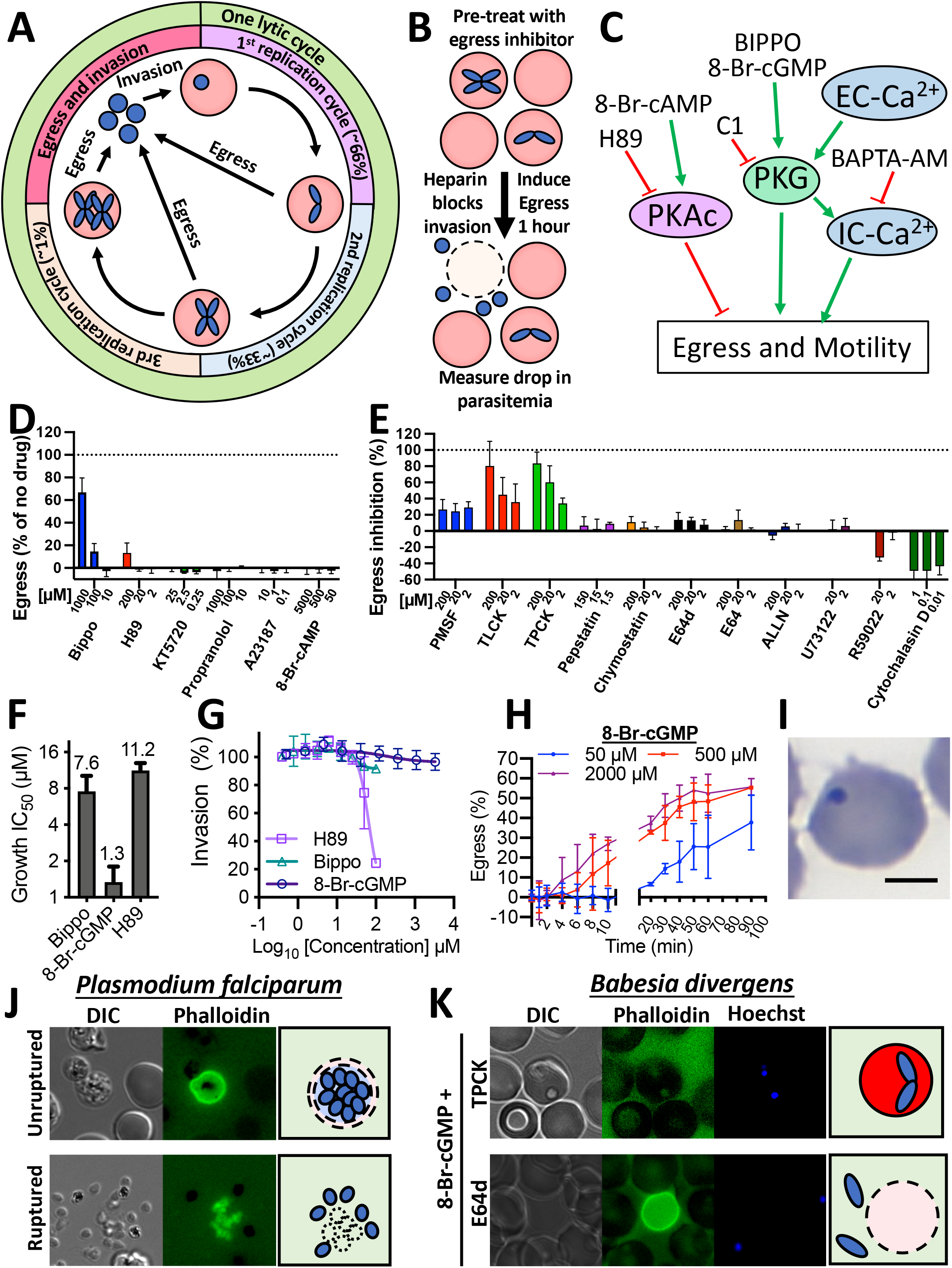
**(A)** Diagram of a lytic cycle of *B. divergens*. One lytic cycle of *B. divergens* can contain multiple replication cycles and includes egress and invasion. The number of replication cycles per lytic cycle is variable. One replication cycle represents a parasite dividing to form two daughters. The percentage shown in brackets represent the approximate percentage of parasites that egress after one, two or more replication cycles. **(B)** Diagram of the flow cytometry-based egress assay. Egress is measured as the reduction in infected cells. **(C)** Model of the egress signaling pathway in *B. divergens* with activators shown with a green arrow, and inhibitors with a red line. **(D)** Percentage of parasites that egress when treated with each compound, as measured by flow cytometry. **(E)** Screen for inhibitors of 8-Br-cGMP mediated induced egress. **(F)** IC_50_ of each compound against proliferation in a 72 hours assay. **(G)** Invasion inhibition by each compound. Isolated merozoites were pre-incubated with the inhibitor for 10 min before adding RBCs. **(H)** Induced egress was measured over time with three concentrations of 8-Br-cGMP. (I) Parasites that have not egressed within 1 hour of 8-Br-cGMP addition become pyknotic. **(J)** *P. falciparum* parasites egressing in the presence of phalloidin, which stains the RBC cytoskeleton. Top. A parasite that has begun egress and permeabilized the RBC. The merozoites remain in the permeabilized RBC. Bottom. Merozoites that have completed egress. The RBC cytoskeleton has fragmented to release the merozoites. (K) Parasites treated with TPCK can not be induced to egress with 8-Br-cGMP. Pre-treatment with E64d does not alter 8-Br-cGMP induced egress.

**Supplementary Figure 2.**
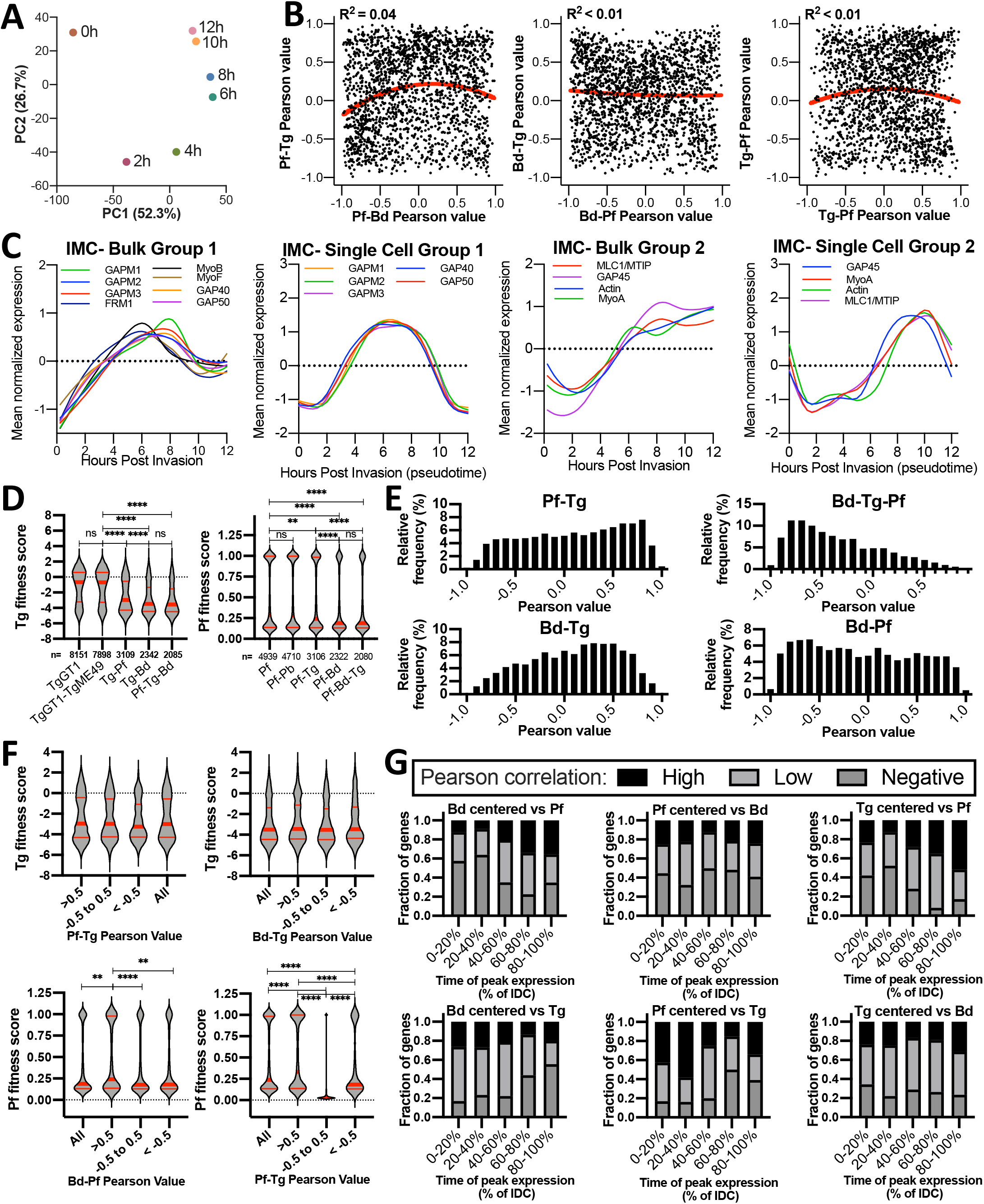
**(A)** PCA plot of each timepoint from the bulk transcriptome demonstrating the cyclical nature of replication. **(B)** Comparison of the Pearson correlation of expression profiles of orthologous genes between species pairs. **(C)** Expression profiles of IMC proteins fall into two clusters. **(D)** Fitness score from *P. falciparum piggyBac* transposon (Zhang et al., 2018) and *T. gondii* CRISPR (Sidik et al., 2016) screens for genes shared between the species on the x-axis. **(E)** The relative frequency of the correlation between expression profiles of two or three species. **(F)** Fitness scores for genes with expression profiles that are positively correlated (>0.5), not correlated (-0.5-0.5) or negatively correlated (< −0.5) between species pairs. **(G)** Pearson correlation between orthologs of species pairs categorized by the time of peak expression of one species (centered spp.). Bd, *B. divergens.* Tg*, T. gondii*. Pf, *P. falciparum*.

**Supplementary Figure 3.**
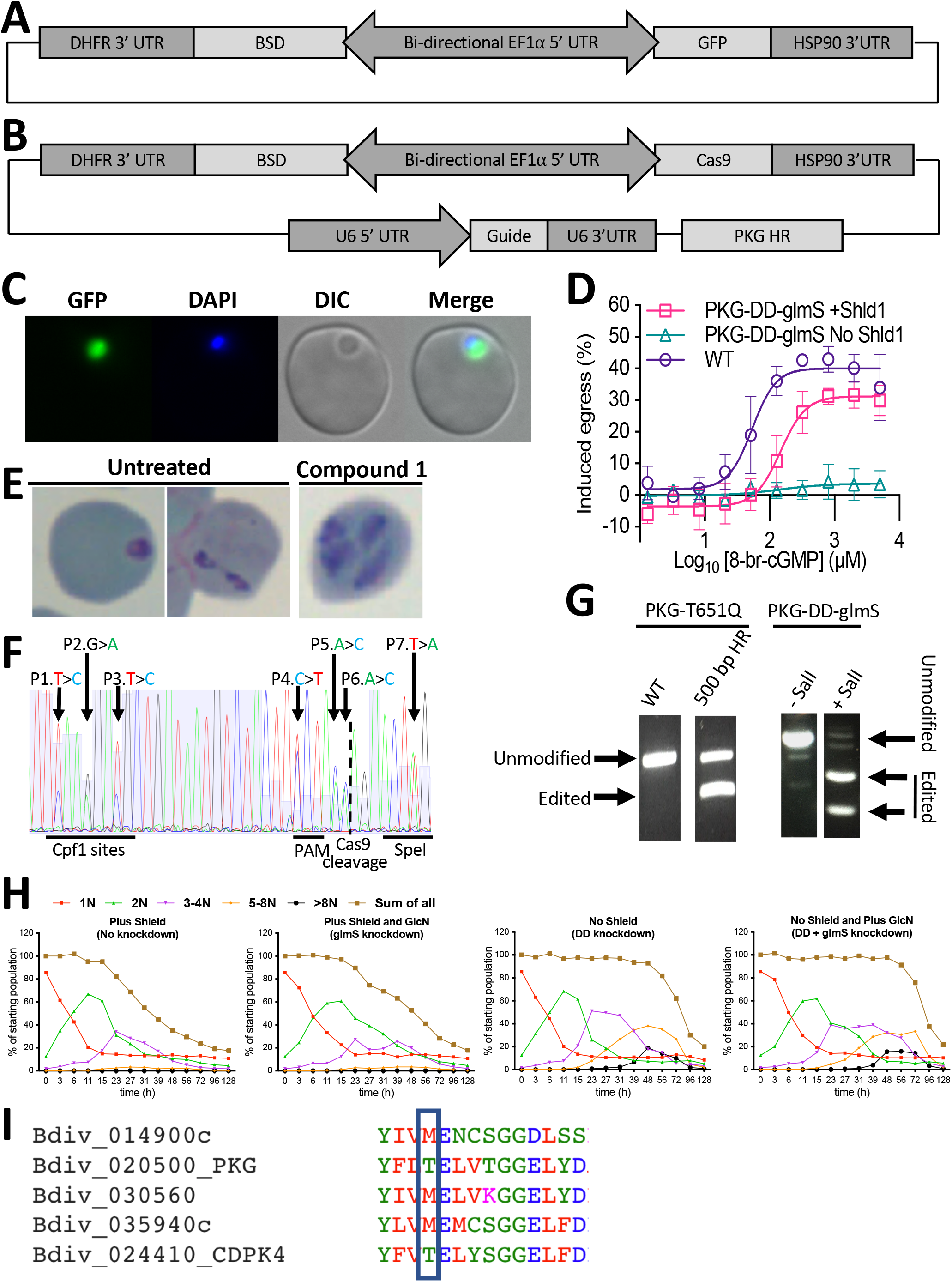
**(A)** Schematic of GFP reporter plasmid. **(B)** Schematic of CRISPR/Cas9 plasmid with a repair template. **(C)** Live cell imaging of *B. divergens* parasites expressing GFP. **(D)** The number of WT or PKG-DD-glmS parasites, with (no Shld1) or without knockdown induced for 24 h, that egress when treated with 8-Br-cGMP. Data is the mean ± SD of three biological experiments performed in technical triplicate **(E)** *B. bovis* parasites treated with C1 display a continued replication phenotype. **(F)** Sanger sequencing of *pkg* in the uncloned parasite population after transfection. Positions 1-7 are the alternative bases in the homology repair template where editing is expected. **(G)** RFLP analysis of the uncloned parasite population after transfection with either a plasmid containing a 500 bp repair template or with a HA-DD-glmS tag flanked by two 500 bp HR regions. (H) The number of parasites per infected RBC over time with PKG knockdown, assessed by flow cytometry. A single representative experiment is shown of three biological repeats in technical triplicate.

**Supplementary Table 1. Orthologs of known apicomplexan egress and invasion genes.**

**Supplementary Table 2.** Putative novel egress and invasion genes in *B. divergens*.

**Supplementary Table 3.** Summary of selection drugs, resistance markers and transfection methods used for *B. divergens*.

**Supplementary Table 4.** Primers used in this study

**Video S1.** 8-Br-cGMP induced egress

**Video S2.** EC no calcium plus 8-Br-cGMP

**Video S3.** 8-Br-cGMP induced egress of Cytochalasin D treated parasites

**Video S4.** IC and EC with calcium plus saponin

**Video S5.** IC and EC without calcium plus saponin

**Video S6.** ASP2 knockdown plus 8-Br-cGMP

**Video S7.** ASP3 knockdown plus 8-Br-cGMP

**Video S8.** ASP3 knockdown plus cytochalasin D plus 8-Br-cGMP

## References

Agarwal, S., Singh, M.K., Garg, S., Chitnis, C.E., and Singh, S. (2013). Ca 2+-mediated exocytosis of subtilisin-like protease 1: a key step in egress of P lasmodium falciparum merozoites. Cellular microbiology. 15(6), 910–921.

Amlabu, E., Mensah-Brown, H., Nyarko, P.B., Akuh, O.-a., Opoku, G., Ilani, P., Oyagbenro, R., Asiedu, K., Aniweh, Y., and Awandare, G.A. (2018). Functional Characterization of Plasmodium falciparum Surface-Related Antigen as a Potential Blood-Stage Vaccine Target. The Journal of infectious diseases. 218(5), 778–790.

Arendse, L.B., Wyllie, S., Chibale, K., and Gilbert, I.H. (2021). Plasmodium kinases as potential drug targets for malaria: challenges and opportunities. ACS Infectious Diseases. 7(3), 518–534.

Armstrong, C.M., and Goldberg, D.E. (2007). An FKBP destabilization domain modulates protein levels in Plasmodium falciparum. Nature methods. 4(12), 1007.

Arumugam, T.U., Takeo, S., Yamasaki, T., Thonkukiatkul, A., Miura, K., Otsuki, H., Zhou, H., Long, C.A., Sattabongkot, J., and Thompson, J. (2011). Discovery of GAMA, a Plasmodium falciparum merozoite micronemal protein, as a novel blood-stage vaccine candidate antigen. Infection and immunity. 79(11), 4523–4532.

Baker, D.A., Matralis, A.N., Osborne, S.A., Large, J.M., and Penzo, M. (2020). Targeting the malaria parasite cGMP-dependent protein kinase to develop new drugs. Frontiers in Microbiology. 3189.

Baker, D.A., Stewart, L.B., Large, J.M., Bowyer, P.W., Ansell, K.H., Jiménez-Díaz, M.B., El Bakkouri, M., Birchall, K., Dechering, K.J., and Bouloc, N.S. (2017). A potent series targeting the malarial cGMP-dependent protein kinase clears infection and blocks transmission. Nature Communications. 8(1), 430.

Behnke, M.S., Wootton, J.C., Lehmann, M.M., Radke, J.B., Lucas, O., Nawas, J., Sibley, L.D., and White, M.W. (2010). Coordinated progression through two subtranscriptomes underlies the tachyzoite cycle of Toxoplasma gondii. PloS one. 5(8), e12354.

Bergmann-Leitner, E.S., Mease, R.M., De La Vega, P., Savranskaya, T., Polhemus, M., Ockenhouse, C., and Angov, E. (2010). Immunization with pre-erythrocytic antigen CelTOS from Plasmodium falciparum elicits cross-species protection against heterologous challenge with Plasmodium berghei. PloS one. 5(8), e12294.

Besteiro, S., Michelin, A., Poncet, J., Dubremetz, J.-F., and Lebrun, M. (2009). Export of a Toxoplasma gondii rhoptry neck protein complex at the host cell membrane to form the moving junction during invasion. PLoS pathog. 5(2), e1000309.

Bhanot, P., Schauer, K., Coppens, I., and Nussenzweig, V. (2005). A surface phospholipase is involved in the migration of Plasmodium sporozoites through cells. Journal of Biological Chemistry. 280(8), 6752–6760.

Bisio, H., Lunghi, M., Brochet, M., and Soldati-Favre, D. (2019). Phosphatidic acid governs natural egress in Toxoplasma gondii via a guanylate cyclase receptor platform. Nature microbiology. 4(3), 420.

Bisio, H., and Soldati-Favre, D. (2019). Signaling cascades governing entry into and exit from host cells by Toxoplasma gondii. Annual review of microbiology. 73, 579–599.

Black, M.W., Arrizabalaga, G., and Boothroyd, J.C. (2000). Ionophore-resistant mutants of Toxoplasma gondii reveal host cell permeabilization as an early event in egress. Molecular and cellular biology. 20(24), 9399–9408.

Bock, R., Jackson, L., De Vos, A., and Jorgensen, W. (2004). Babesiosis of cattle. Parasitology. 129(S1), S247–S269.

Bozdech, Z., Llinás, M., Pulliam, B.L., Wong, E.D., Zhu, J., and DeRisi, J.L. (2003). The transcriptome of the intraerythrocytic developmental cycle of Plasmodium falciparum. PLoS Biol. 1(1), e5.

Brown, K.M., Long, S., and Sibley, L.D. (2017). Plasma Membrane Association by N-Acylation Governs PKG Function in Toxoplasma gondii. mBio. 8(3), e00375–00317.

Brown, K.M., Lourido, S., and Sibley, L.D. (2016). Serum albumin stimulates protein kinase G-dependent microneme secretion in Toxoplasma gondii. Journal of Biological Chemistry. 291(18), 9554–9565.

Bullen, H.E., Jia, Y., Yamaryo-Botté, Y., Bisio, H., Zhang, O., Jemelin, N.K., Marq, J.-B., Carruthers, V., Botté, C.Y., and Soldati-Favre, D. (2016). Phosphatidic acid-mediated signaling regulates microneme secretion in Toxoplasma. Cell host & microbe. 19(3), 349–360.

Caldas, L.A., and De Souza, W. (2018). A window to Toxoplasma gondii egress. Pathogens. 7(3), 69.

Chandramohanadas, R., Davis, P.H., Beiting, D.P., Harbut, M.B., Darling, C., Velmourougane, G., Lee, M.Y., Greer, P.A., Roos, D.S., and Greenbaum, D.C. (2009). Apicomplexan parasites co-opt host calpains to facilitate their escape from infected cells. science. 324(5928), 794–797.

Chappell, L., Ross, P., Orchard, L., Russell, T.J., Otto, T.D., Berriman, M., Rayner, J.C., and Llinás, M. (2020). Refining the transcriptome of the human malaria parasite Plasmodium falciparum using amplification-free RNA-seq. BMC genomics. 21(1), 1–19.

Collins, C.R., Hackett, F., Strath, M., Penzo, M., Withers-Martinez, C., Baker, D.A., and Blackman, M.J. (2013). Malaria parasite cGMP-dependent protein kinase regulates blood stage merozoite secretory organelle discharge and egress. PLoS Pathog. 9(5), e1003344.

Cursino-Santos, J.R., Singh, M., Pham, P., Rodriguez, M., and Lobo, C.A. (2016). Babesia divergens builds a complex population structure composed of specific ratios of infected cells to ensure a prompt response to changing environmental conditions. Cellular microbiology. 18(6), 859–874.

Das, S., Lemgruber, L., Tay, C.L., Baum, J., and Meissner, M. (2017). Multiple essential functions of Plasmodium falciparum actin-1 during malaria blood-stage development. BMC biology. 15(1), 1–16.

Deligianni, E., de Monerri, N.C.S., McMillan, P.J., Bertuccini, L., Superti, F., Manola, M., Spanos, L., Louis, C., Blackman, M.J., and Tilley, L. (2018). Essential role of Plasmodium perforin-like protein 4 in ookinete midgut passage. PloS one. 13(8), e0201651.

Deligianni, E., Morgan, R.N., Bertuccini, L., Wirth, C.C., Silmon de Monerri, N.C., Spanos, L., Blackman, M.J., Louis, C., Pradel, G., and Siden-Kiamos, I. (2013). A perforin-like protein mediates disruption of the erythrocyte membrane during egress of P lasmodium berghei male gametocytes. Cellular microbiology. 15(8), 1438–1455.

Diaz, C.A., Allocco, J., Powles, M.A., Yeung, L., Donald, R.G.K., Anderson, J.W., and Liberator, P.A. (2006). Characterization of Plasmodium falciparum cGMP-dependent protein kinase (PfPKG): Antiparasitic activity of a PKG inhibitor. Molecular and Biochemical Parasitology. 146(1), 78–88. DOI: https://doi.org/10.1016/j.molbiopara.2005.10.020.

Dogga, S.K., Mukherjee, B., Jacot, D., Kockmann, T., Molino, L., Hammoudi, P.-M., Hartkoorn, R.C., Hehl, A.B., and Soldati-Favre, D. (2017). A druggable secretory protein maturase of Toxoplasma essential for invasion and egress. Elife. 6, e27480.

Dominicus, C., Nofal, S.D., Broncel, M., Katris, N.J., Flynn, H.R., Arrizabalaga, G., Botté, C.Y., Invergo, B.M., and Treeck, M. (2021). A positive feedback loop mediates crosstalk between calcium, cyclic nucleotide and lipid signalling in Toxoplasma gondii. bioRxiv.

Donald, R.G., Zhong, T., Wiersma, H., Nare, B., Yao, D., Lee, A., Allocco, J., and Liberator, P.A. (2006). Anticoccidial kinase inhibitors: identification of protein kinase targets secondary to cGMP-dependent protein kinase. Molecular and biochemical parasitology. 149(1), 86–98.

Dunkley, T.P., Watson, R., Griffin, J.L., Dupree, P., and Lilley, K.S. (2004). Localization of organelle proteins by isotope tagging (LOPIT). Molecular & Cellular Proteomics. 3(11), 1128–1134.

Dvorin, J.D., Bei, A.K., Coleman, B.I., and Duraisingh, M.T. (2010). Functional diversification between two related Plasmodium falciparum merozoite invasion ligands is determined by changes in the cytoplasmic domain. Mol Microbiol. 75(4), 990–1006. Published online 2010/05/22 DOI: 10.1111/j.1365-2958.2009.07040.x.

Favuzza, P., de Lera Ruiz, M., Thompson, J.K., Triglia, T., Ngo, A., Steel, R.W., Vavrek, M., Christensen, J., Healer, J., and Boyce, C. (2020). Dual plasmepsin-targeting antimalarial agents disrupt multiple stages of the malaria parasite life cycle. Cell host & microbe. 27(4), 642–658. e612.

Flueck, C., Drought, L.G., Jones, A., Patel, A., Perrin, A.J., Walker, E.M., Nofal, S.D., Snijders, A.P., Blackman, M.J., and Baker, D.A. (2019). Phosphodiesterase beta is the master regulator of cAMP signalling during malaria parasite invasion. PLoS biology. 17(2), e3000154.

Gaji, R.Y., Behnke, M.S., Lehmann, M.M., White, M.W., and Carruthers, V.B. (2011). Cell cycle-dependent, intercellular transmission of Toxoplasma gondii is accompanied by marked changes in parasite gene expression. Molecular microbiology. 79(1), 192–204.

Ghai, M., Dutta, S., Hall, T., Freilich, D., and Ockenhouse, C.F. (2002). Identification, expression, and functional characterization of MAEBL, a sporozoite and asexual blood stage chimeric erythrocyte-binding protein of Plasmodium falciparum. Molecular and biochemical parasitology. 123(1), 35–45.

Ghosh, S., Chisholm, S.A., Dans, M., Lakkavaram, A., Kennedy, K., Ralph, S.A., Counihan, N.A., and de Koning-Ward, T.F. (2018). The cysteine protease dipeptidyl aminopeptidase 3 does not contribute to egress of Plasmodium falciparum from host red blood cells. PLoS One. 13(3), e0193538.

Gilson, P.R., and Crabb, B.S. (2009). Morphology and kinetics of the three distinct phases of red blood cell invasion by Plasmodium falciparum merozoites. International journal for parasitology. 39(1), 91–96.

Glushakova, S., Beck, J.R., Garten, M., Busse, B.L., Nasamu, A.S., Tenkova-Heuser, T., Heuser, J., Goldberg, D.E., and Zimmerberg, J. (2018). Rounding precedes rupture and breakdown of vacuolar membranes minutes before malaria parasite egress from erythrocytes. Cell Microbiol. 20(10), e12868. Published online 2018/06/15 DOI: 10.1111/cmi.12868.

Glushakova, S., Humphrey, G., Leikina, E., Balaban, A., Miller, J., and Zimmerberg, J. (2010). New stages in the program of malaria parasite egress imaged in normal and sickle erythrocytes. Current Biology. 20(12), 1117–1121.

González, L.M., Estrada, K., Grande, R., Jiménez-Jacinto, V., Vega-Alvarado, L., Sevilla, E., de la Barrera, J., Cuesta, I., Zaballos, Á., and Bautista, J.M. (2019). Comparative and functional genomics of the protozoan parasite Babesia divergens highlighting the invasion and egress processes. PLoS neglected tropical diseases. 13(8), e0007680.

Gubbels, M.-J., Keroack, C.D., Dangoudoubiyam, S., Worliczek, H.L., Paul, A.S., Bauwens, C., Elsworth, B., Engelberg, K., Howe, D.K., and Coppens, I. (2020). Fussing about fission: defining variety among mainstream and exotic apicomplexan cell division modes. Frontiers in Cellular and Infection Microbiology. 10, 269.

Gurnett, A.M., Liberator, P.A., Dulski, P.M., Salowe, S.P., Donald, R.G., Anderson, J.W., Wiltsie, J., Diaz, C.A., Harris, G., and Chang, B. (2002). Purification and molecular characterization of cGMP-dependent protein kinase from apicomplexan parasites a novel chemotherapeutic target. Journal of Biological Chemistry. 277(18), 15913–15922.

Hale, V.L., Watermeyer, J.M., Hackett, F., Vizcay-Barrena, G., van Ooij, C., Thomas, J.A., Spink, M.C., Harkiolaki, M., Duke, E., Fleck, R.A., et al. (2017). Parasitophorous vacuole poration precedes its rupture and rapid host erythrocyte cytoskeleton collapse in Plasmodium falciparum egress. Proc Natl Acad Sci U S A. 114(13), 3439–3444. Published online 2017/03/16 DOI: 10.1073/pnas.1619441114.

Hassan, M.A., Vasquez, J.J., Guo-Liang, C., Meissner, M., and Nicolai Siegel, T. (2017). Comparative ribosome profiling uncovers a dominant role for translational control in Toxoplasma gondii. BMC genomics. 18(1), 1–12.

Hehl, A.B., Basso, W.U., Lippuner, C., Ramakrishnan, C., Okoniewski, M., Walker, R.A., Grigg, M.E., Smith, N.C., and Deplazes, P. (2015). Asexual expansion of Toxoplasma gondii merozoites is distinct from tachyzoites and entails expression of non-overlapping gene families to attach, invade, and replicate within feline enterocytes. BMC genomics. 16(1), 1–16.

Howard, B.L., Harvey, K.L., Stewart, R.J., Azevedo, M.F., Crabb, B.S., Jennings, I.G., Sanders, P.R., Manallack, D.T., Thompson, P.E., and Tonkin, C.J. (2015). Identification of potent phosphodiesterase inhibitors that demonstrate cyclic nucleotide-dependent functions in apicomplexan parasites. ACS chemical biology. 10(4), 1145–1154.

Hui, R., El Bakkouri, M., and Sibley, L.D. (2015). Designing selective inhibitors for calcium-dependent protein kinases in apicomplexans. Trends in pharmacological sciences. 36(7), 452–460.

Jia, Y., Marq, J.B., Bisio, H., Jacot, D., Mueller, C., Yu, L., Choudhary, J., Brochet, M., and Soldati-Favre, D. (2017). Crosstalk between PKA and PKG controls pH-dependent host cell egress of *Toxoplasma gondii*. The EMBO Journal. DOI: 10.15252/embj.201796794.

Kafsack, B.F., Pena, J.D., Coppens, I., Ravindran, S., Boothroyd, J.C., and Carruthers, V.B. (2009). Rapid membrane disruption by a perforin-like protein facilitates parasite exit from host cells. Science. 323(5913), 530–533.

Lal, K., Prieto, J.H., Bromley, E., Sanderson, S.J., Yates III, J.R., Wastling, J.M., Tomley, F.M., and Sinden, R.E. (2009). Characterisation of Plasmodium invasive organelles; an ookinete microneme proteome. Proteomics. 9(5), 1142–1151.

Lamarque, M., Besteiro, S., Papoin, J., Roques, M., Vulliez-Le Normand, B., Morlon-Guyot, J., Dubremetz, J.-F., Fauquenoy, S., Tomavo, S., and Faber, B.W. (2011). The RON2-AMA1 interaction is a critical step in moving junction-dependent invasion by apicomplexan parasites. PLoS pathogens. 7(2), e1001276.

Lau, A.O., Pedroni, M.J., and Bhanot, P. (2013). Target specific-trisubstituted pyrrole inhibits Babesia bovis erythrocytic growth. Experimental parasitology. 133(3), 365–368.

Lehmann, C., Tan, M.S.Y., de Vries, L.E., Russo, I., Sanchez, M.I., Goldberg, D.E., and Deu, E. (2018). Plasmodium falciparum dipeptidyl aminopeptidase 3 activity is important for efficient erythrocyte invasion by the malaria parasite. PLoS pathogens. 14(5), e1007031.

Lyon, J., and Haynes, J. (1986). Plasmodium falciparum antigens synthesized by schizonts and stabilized at the merozoite surface when schizonts mature in the presence of protease inhibitors. The Journal of Immunology. 136(6), 2245–2251.

McRobert, L., Taylor, C.J., Deng, W., Fivelman, Q.L., Cummings, R.M., Polley, S.D., Billker, O., and Baker, D.A. (2008). Gametogenesis in Malaria Parasites Is Mediated by the cGMP-Dependent Protein Kinase. PLOS Biology. 6(6), e139. DOI: 10.1371/journal.pbio.0060139.

Moss, W.J., Patterson, C.E., Jochmans, A.K., and Brown, K.M. (2021). Functional Analysis of the Expanded Phosphodiesterase Gene Family in Toxoplasma gondii Tachyzoites. bioRxiv.

Mossaad, E., Asada, M., Nakatani, D., Inoue, N., Yokoyama, N., Kaneko, O., and Kawazu, S.-i. (2015). Calcium ions are involved in egress of Babesia bovis merozoites from bovine erythrocytes. Journal of Veterinary Medical Science. 77(1), 53–58.

Moudy, R., Manning, T.J., and Beckers, C.J. (2001). The loss of cytoplasmic potassium upon host cell breakdown triggers egress of Toxoplasma gondii. Journal of Biological Chemistry. 276(44), 41492–41501.

Nasamu, A.S., Glushakova, S., Russo, I., Vaupel, B., Oksman, A., Kim, A.S., Fremont, D.H., Tolia, N., Beck, J.R., and Meyers, M.J. (2017). Plasmepsins IX and X are essential and druggable mediators of malaria parasite egress and invasion. Science. 358(6362), 518–522.

Otto, T.D., Wilinski, D., Assefa, S., Keane, T.M., Sarry, L.R., Böhme, U., Lemieux, J., Barrell, B., Pain, A., and Berriman, M. (2010). New insights into the blood-stage transcriptome of Plasmodium falciparum using RNA-Seq. Molecular microbiology. 76(1), 12–24.

Paoletta, M.S., Laughery, J.M., Arias, L.S.L., Ortiz, J.M.J., Montenegro, V.N., Petrigh, R., Ueti, M.W., Suarez, C.E., Farber, M.D., and Wilkowsky, S.E. (2021). The key to egress? Babesia bovis perforin-like protein 1 (PLP1) with hemolytic capacity is required for blood stage replication and is involved in the exit of the parasite from the host cell. International journal for parasitology. 51(8), 643–658.

Patel, A., Perrin, A.J., Flynn, H.R., Bisson, C., Withers-Martinez, C., Treeck, M., Flueck, C., Nicastro, G., Martin, S.R., and Ramos, A. (2019). Cyclic AMP signalling controls key components of malaria parasite host cell invasion machinery. PLoS biology. 17(5), e3000264.

Paul, A.S., Miliu, A., Paulo, J.A., Goldberg, J.M., Bonilla, A.M., Berry, L., Seveno, M., Braun-Breton, C., Kosber, A.L., Elsworth, B., et al. (2020). Co-option of Plasmodium falciparum PP1 for egress from host erythrocytes. Nature Communications. 11(1), 3532. DOI: 10.1038/s41467-020-17306-1.

Pedroni, M.J., Sondgeroth, K.S., Gallego-Lopez, G.M., Echaide, I., and Lau, A.O. (2013). Comparative transcriptome analysis of geographically distinct virulent and attenuated Babesia bovis strains reveals similar gene expression changes through attenuation. BMC genomics. 14(1), 763.

Peloakgosi-Shikwambani, K., 2018. Analysis of Babesia rossi transcriptome in dogs diagnosed with canine babesiosis.

Perrin, A.J., Collins, C.R., Russell, M.R., Collinson, L.M., Baker, D.A., and Blackman, M.J. (2018). The actinomyosin motor drives malaria parasite red blood cell invasion but not egress. MBio. 9(4), e00905–00918.

Picelli, S., Björklund, Å.K., Faridani, O.R., Sagasser, S., Winberg, G., and Sandberg, R. (2013). Smart-seq2 for sensitive full-length transcriptome profiling in single cells. Nature methods. 10(11), 1096–1098.

Pillai, A.D., Addo, R., Sharma, P., Nguitragool, W., Srinivasan, P., and Desai, S.A. (2013). Malaria parasites tolerate a broad range of ionic environments and do not require host cation remodelling. Molecular microbiology. 88(1), 20–34.

Pino, P., Caldelari, R., Mukherjee, B., Vahokoski, J., Klages, N., Maco, B., Collins, C.R., Blackman, M.J., Kursula, I., and Heussler, V. (2017). A multistage antimalarial targets the plasmepsins IX and X essential for invasion and egress. Science. 358(6362), 522–528.

Prommana, P., Uthaipibull, C., Wongsombat, C., Kamchonwongpaisan, S., Yuthavong, Y., Knuepfer, E., Holder, A.A., and Shaw, P.J. (2013). Inducible knockdown of Plasmodium gene expression using the glmS ribozyme. PloS one. 8(8), e73783.

Rezvani, Y., Keroack, C.D., Elsworth, B., Arriojas, A., Gubbels, M.-J., Duraisingh, M.T., and Zarringhalam, K. (2022). Comparative single-cell transcriptional atlases of *Babesia* species reveal conserved and species-specific expression profiles. bioRxiv. 2022.2002.2011.480160. DOI: 10.1101/2022.02.11.480160.

Rossouw, I., Maritz-Olivier, C., Niemand, J., Van Biljon, R., Smit, A., Olivier, N.A., and Birkholtz, L.-M. (2015). Morphological and molecular descriptors of the developmental cycle of Babesia divergens parasites in human erythrocytes. PLoS Negl Trop Dis. 9(5), e0003711.

Rupp, I., Bosse, R., Schirmeister, T., and Pradel, G. (2008). Effect of protease inhibitors on exflagellation in Plasmodium falciparum. Molecular and biochemical parasitology. 158(2), 208–212.

Salmon, B.L., Oksman, A., and Goldberg, D.E. (2001). Malaria parasite exit from the host erythrocyte: a two-step process requiring extraerythrocytic proteolysis. Proceedings of the National Academy of Sciences. 98(1), 271–276.

Sassmannshausen, J., Pradel, G., and Bennink, S. (2020). Perforin-like proteins of apicomplexan parasites. Frontiers in Cellular and Infection Microbiology. 10.

Schultz, A.J., and Carruthers, V.B. (2018). Toxoplasma gondii LCAT Primarily Contributes to Tachyzoite Egress. mSphere. 3(1), e00073–00018.

Sidik, S.M., Huet, D., Ganesan, S.M., Huynh, M.-H., Wang, T., Nasamu, A.S., Thiru, P., Saeij, J.P., Carruthers, V.B., and Niles, J.C. (2016). A genome-wide CRISPR screen in Toxoplasma identifies essential apicomplexan genes. Cell. 166(6), 1423–1435. e1412.

Silva, J.C., Cornillot, E., McCracken, C., Usmani-Brown, S., Dwivedi, A., Ifeonu, O.O., Crabtree, J., Gotia, H.T., Virji, A.Z., and Reynes, C. (2016). Genome-wide diversity and gene expression profiling of Babesia microti isolates identify polymorphic genes that mediate host-pathogen interactions. Scientific reports. 6, 35284.

Singh, S., Alam, M.M., Pal-Bhowmick, I., Brzostowski, J.A., and Chitnis, C.E. (2010). Distinct external signals trigger sequential release of apical organelles during erythrocyte invasion by malaria parasites. PLoS Pathog. 6(2), e1000746.

Singh, S., and Chitnis, C.E. (2017). Molecular signaling involved in entry and exit of malaria parasites from host erythrocytes. Cold Spring Harbor Perspectives in Medicine. 7(10), a026815.

Suarez, C., Bishop, R., Alzan, H., Poole, W., and Cooke, B. (2017). Advances in the application of genetic manipulation methods to apicomplexan parasites. International journal for parasitology. 47(12), 701–710.

Suárez-Cortés, P., Sharma, V., Bertuccini, L., Costa, G., Bannerman, N.-L., Sannella, A.R., Williamson, K., Klemba, M., Levashina, E.A., and Lasonder, E. (2016). Comparative proteomics and functional analysis reveal a role of Plasmodium falciparum osmiophilic bodies in malaria parasite transmission. Molecular & Cellular Proteomics. 15(10), 3243–3255.

Swearingen, K.E., Lindner, S.E., Shi, L., Shears, M.J., Harupa, A., Hopp, C.S., Vaughan, A.M., Springer, T.A., Moritz, R.L., and Kappe, S.H. (2016). Interrogating the Plasmodium sporozoite surface: identification of surface-exposed proteins and demonstration of glycosylation on CSP and TRAP by mass spectrometry-based proteomics. PLoS pathogens. 12(4), e1005606.

Tan, M.S.Y., and Blackman, M.J. (2021). Malaria parasite egress at a glance. Journal of cell science. 134(5), jcs257345.

Terkawi, M.A., Ratthanophart, J., Salama, A., AbouLaila, M., Asada, M., Ueno, A., Alhasan, H., Guswanto, A., Masatani, T., and Yokoyama, N. (2013). Molecular characterization of a new Babesia bovis thrombospondin-related anonymous protein (BbTRAP2). PLoS One. 8(12), e83305.

Thomas, J.A., Tan, M.S.Y., Bisson, C., Borg, A., Umrekar, T.R., Hackett, F., Hale, V.L., Vizcay-Barrena, G., Fleck, R.A., Snijders, A.P., et al. (2018). A protease cascade regulates release of the human malaria parasite Plasmodium falciparum from host red blood cells. Nat Microbiol. 3(4), 447–455. Published online 2018/02/21 DOI: 10.1038/s41564-018-0111-0.

Treeck, M., Zacherl, S., Herrmann, S., Cabrera, A., Kono, M., Struck, N.S., Engelberg, K., Haase, S., Frischknecht, F., and Miura, K. (2009). Functional analysis of the leading malaria vaccine candidate AMA-1 reveals an essential role for the cytoplasmic domain in the invasion process. PLoS pathogens. 5(3), e1000322.

Uboldi, A.D., Wilde, M.-L., McRae, E.A., Stewart, R.J., Dagley, L.F., Yang, L., Katris, N.J., Hapuarachchi, S.V., Coffey, M.J., and Lehane, A.M. (2018). Protein kinase A negatively regulates Ca2+ signalling in Toxoplasma gondii. PLoS biology. 16(9), e2005642.

Ueti, M.W., Johnson, W.C., Kappmeyer, L.S., Herndon, D.R., Mousel, M.R., Reif, K.E., Taus, N.S., Ifeonu, O.O., Silva, J.C., and Suarez, C.E. (2020). Transcriptome dataset of Babesia bovis life stages within vertebrate and invertebrate hosts. Data in Brief. 106533.

Van Voorhis, W.C., Doggett, J.S., Parsons, M., Hulverson, M.A., Choi, R., Arnold, S.L., Riggs, M.W., Hemphill, A., Howe, D.K., and Mealey, R.H. (2017). Extended-spectrum antiprotozoal bumped kinase inhibitors: a review. Experimental parasitology. 180, 71–83.

Vella, S.A., Moore, C.A., Li, Z.-H., Triana, M.A.H., Potapenko, E., and Moreno, S.N. (2021). The role of potassium and host calcium signaling in Toxoplasma gondii egress. Cell Calcium. 94, 102337.

Wasserman, M., Alarcón, C., and Mendoza, P.M. (1982). Effects of Ca++ depletion on the asexual cell cycle of Plasmodium falciparum. The American journal of tropical medicine and hygiene. 31(4), 711–717.

Weiss, G.E., Gilson, P.R., Taechalertpaisarn, T., Tham, W.-H., de Jong, N.W., Harvey, K.L., Fowkes, F.J., Barlow, P.N., Rayner, J.C., and Wright, G.J. (2015a). Revealing the sequence and resulting cellular morphology of receptor-ligand interactions during Plasmodium falciparum invasion of erythrocytes. PLoS pathogens. 11(2), e1004670.

Weiss, G.E., Gilson, P.R., Taechalertpaisarn, T., Tham, W.-H., de Jong, N.W., Harvey, K.L., Fowkes, F.J., Barlow, P.N., Rayner, J.C., and Wright, G.J. (2015b). Revealing the sequence and resulting cellular morphology of receptor-ligand interactions during Plasmodium falciparum invasion of erythrocytes. PLoS Pathog. 11(2), e1004670.

Wilde, M.-L., Triglia, T., Marapana, D., Thompson, J.K., Kouzmitchev, A.A., Bullen, H.E., Gilson, P.R., Cowman, A.F., and Tonkin, C.J. (2019). Protein kinase A is essential for invasion of Plasmodium falciparum into human erythrocytes. mBio. 10(5), e01972–01919.

Wirth, C.C., Glushakova, S., Scheuermayer, M., Repnik, U., Garg, S., Schaack, D., Kachman, M.M., Weißbach, T., Zimmerberg, J., and Dandekar, T. (2014). Perforin-like protein PPLP 2 permeabilizes the red blood cell membrane during egress of P lasmodium falciparum gametocytes. Cellular microbiology. 16(5), 709–733.

Wright, I., Casu, R., Commins, M., Dalrymple, B., Gale, K., Goodger, B., Riddles, P., Waltisbuhl, D., Abetz, I., and Berrie, D. (1992). The development of a recombinant Babesia vaccine. Veterinary parasitology. 44(1-2), 3–13.

Yap, A., Azevedo, M.F., Gilson, P.R., Weiss, G.E., O’Neill, M.T., Wilson, D.W., Crabb, B.S., and Cowman, A.F. (2014). Conditional expression of apical membrane antigen 1 in P lasmodium falciparum shows it is required for erythrocyte invasion by merozoites. Cellular microbiology. 16(5), 642–656.

Young, J.A., Johnson, J.R., Benner, C., Yan, S.F., Chen, K., Le Roch, K.G., Zhou, Y., and Winzeler, E.A. (2008). In silico discovery of transcription regulatory elements in Plasmodium falciparum. BMC genomics. 9(1), 1–21.

Zhang, M., Wang, C., Otto, T.D., Oberstaller, J., Liao, X., Adapa, S.R., Udenze, K., Bronner, I.F., Casandra, D., Mayho, M., et al. (2018). Uncovering the essential genes of the human malaria parasite *Plasmodium falciparum* by saturation mutagenesis. Science. 360(6388). DOI: 10.1126/science.aap7847.

Zhou, J., Fukumoto, S., Jia, H., Yokoyama, N., Zhang, G., Fujisaki, K., Lin, J., and Xuan, X. (2006). Characterization of the Babesia gibsoni P18 as a homologue of thrombospondin related adhesive protein. Molecular and biochemical parasitology. 148(2), 190–198.

