## Supplementary Table 3. Summary of selection drugs, resistance markers and transfection methods used for B. divergens. for "*Babesia divergens* egress from host cells is orchestrated by essential and druggable kinases and proteases"

| **Selection Markers** | | | | |
| --- | --- | --- | --- | --- |
| **Resistance gene** | **Selection drug** | **IC_50_** | **Concentration used for selection** | **Positive transfection** |
| hDHFR | Pyrimethamine | >500 nM | N/A | N/A |
|  | WR99210 | 43 nM | 100 nM | No |
| BSD | Blasticidin-S | 12 µg/ml | 20 µg/ml | **Yes** |
| PAC | Puromycin | 0.22 µg/ml | 0.5 µg/ml | **Yes (less reliable)** |
| yDHODH | DSM-1 | >100 µM | N/A | N/A |
|  | Atovaquone | 24 nM | 100 nM | No (spontaneous resistance) |
| HYG | Hygromycin B | >500 µg/ml | N/A | N/A |
| **Transfection Methods** | | | | |
| **Transfection System** | **Protocol** | **DNA transfected** | **Parasites transfected** | **Time to reach 1% parasitemia** |
| Bio-rad gene pulser II | 310 V, 950 µF,  ∞ ohms. Cytomix | 100 µg | 200 µl packed iRBCs,  ~20-30% parasitemia | 21 days |
| Amaxa 4D nucleofection | FP158 / P3 solution  100 µl | 10 µg | Free merozoites  From 1.2 ml packed iRBCs, ~20-30% parasitemia | 9 days |
| Amaxa 4D nucleofection | FP158 / P3 solution  100 µl | 10 µg | 20 µl packed iRBCs ~20-30% parasitemia | 14 days |
