## Supplementary material for "*Babesia divergens* egress from host cells is orchestrated by essential and druggable kinases and proteases": Table S4

| BE-8 | ATGACTCGAGGCTAGCGTTTGGCACTGTTGCTCC | Fw Bd. EF1 5’UTR | XhoI-NheI |
| --- | --- | --- | --- |
| BE-9 | ATGAGGATCCGATTACAAGGTACCATTAATCTGCAAAGG | Rv Bd. EF1 5’UTR | BamHI |
| BE-21 | ATGAAAGCTTGTTATACCGCTTTAAGCTGG | Fw Bd DHFR 3’UTR | HindIII |
| BE-22 | ATGAGAATTCGTTGCGATATATATTGACTCTGTTTC | Rv Bd DHFR 3’UTR | EcoRI |
| BE-32 | ATGAACTAGTATATGTAATATAAAGGCTATGTGTATGCATC | Fw Bd Hsp90 3’UTR | SpeI |
| BE-33 | ATGAGCGGCCGCCGCTGATGTGGCACTAGC | Rv Bd Hsp90 3’UTR | NotI |
| BE_124 | gcagattaatggtaccttgtaatcctcgagATGGACTATAAGGACCACGACGG | Fw Cas9 for Bd-GFP-BSD plasmid | Gibson/XhoI |
| BE_125 | acacatagcctttatattacatatactagtTTACTTTTTCTTTTTTGCCTGGCCG | Rv Cas9 for Bd-GFP-BSD plasmid | Gibson/SpeI |
| BE_154 | ggttCTCCAGTCACAAGTTCTGTT | Fw Bd cas9 guide PKG T651Q |  |
| BE_155 | aaacAACAGAACTTGTGACTGGAG | Rv Bd cas9 guide PKG T651Q |  |
| BE_179 | GCACGGTTGGTTTTTTAATACACCG | Fw BdPKG gatekeeper outside HR (check integration) |  |
| BE_180 | ACCAAGTATGATTTCAGGTGCC | Rv BdPKG gatekeeper outside HR (check integration) |  |
| BE_209 | accggtTACCCTTACGATGTTCCTGACTATGC | Fw HA for Bd CRISPR tagging |  |
| BE_212 | gtcgacAGATCATGTGATTTCTCTTTGTTCAAGG | Rv glmS for Bd CRISPR tagging |  |
| BE_213 | tgggatatattagccgtatcGTTTGGCCGATGTATGGTCCTTTGG | Fw PKG gene end for CRISPR tagging. |  |
| BE_215 | GAAATCACATGATCTgtcgacACATAGTTATGAAGCAACTGATTGTGAGC | Fw PKG 3’UTR HR. CRISPR tagging. |  |
| BE_216 | ctcactatagaattcttaattaaCAAACGTGACAGATGGCATCTGAACC | Rv PKG 3’UTR HR. CRISPR tagging. |  |
| BE_221 | ggttAGACCTCACAGACGAAGATT | Fw Cas9 Guide for PKG tagging |  |
| BE_222 | aaacAATCTTCGTCTGTGAGGTCT | Rv Cas9 Guide for PKG tagging |  |
| BE_273 | GCATATAGCACATAATGCGTCTCAGG | Fw Check PKG tag outside HR |  |
| BE_274 | TGCTGGTTGAGTCGCTTGTCC | Rv Check PKG tag outside HR |  |
| BE_366 | TAACGGTCCCGAAAGTTCCTATACC | Rv altered chimeric gDNA for Babesia plasmids | AvrII-gibson |
| BE_367 | ACAAGTATCGATACAGGGAGGTTggGTCTTCgaGAAGACctgtttGagagctaTGCTGGAACAGCAtagcaagttCaaataaggctagtcc | Fw altered chimeric gDNA for Babesia plasmids | Bbs1-gibson |
| BE_494 | tgggatatattagccgtatcctaggCGGATAATGCTCCAGCAGTCG | Fw HR1 BdPLP1 | Gibson- AvrII |
| BE_495 | acatcgtaagggtaaccggtTTGTTTGCGGTCTGATAGTTCTGGT | Rv HR1 BdPLP1 | Gibson to 5’HA |
| BE_496 | gaaatcacatgatctgtcgacTCGCACTGTCATGTACCATAGATTTAAACG | Fw HR2 BdPLP1 | Gibson to glmS 3’ |
| BE_497 | ctcactatagaattcttaattaaTGATGTGACTCTTGCACATACAGC | Rv HR2 BdPLP1 | Gibson-PacI |
| BE_500 | tgggatatattagccgtatcctaggcagAATGAGATAGCGGTATCCTGC | Fw HR1 BdPLP3 | Gibson- AvrII |
| BE_501 | acatcgtaagggtaaccggtTGTTTTAATGTTGTTaGTgTCGGCAC | Rv HR1 BdPLP3 | Gibson to 5’HA |
| BE_502 | gaaatcacatgatctgtcgacCATCCAAAAGTATCACAATCACGACG | Fw HR2 BdPLP3 | Gibson to glmS 3’ |
| BE_503 | ctcactatagaattcttaattaaACAGCAGAAGACTAAGTGTTCATACG | Rv HR2 BdPLP3 | Gibson-PacI |
| BE_506 | tgggatatattagccgtatcctaggCCTTTGAAGTGCTAAACATGGAGG | Fw HR1 BdCDPK5 | Gibson- AvrII |
| BE_507 | acatcgtaagggtaaccggtGTCTAATGTGGCTACATCTCCGG | Rv HR1 BdCDPK5 | Gibson to 5’HA |
| BE_508 | gaaatcacatgatctgtcgacCTCATTCAAACTAAGATGTACACTACGTTTACAC | Fw HR2 BdCDPK5 | Gibson to glmS 3’ |
| BE_509 | ctcactatagaattcttaattaaTGCATTGGGTTCCCTTTTCCC | Rv HR2 BdCDPK5 | Gibson-PacI |
| BE_512 | tgggatatattagccgtatcctaggTCACGCAGAAACTCTCACAAGC | Fw HR1 BdCDPK4 | Gibson- AvrII |
| BE_513 | acatcgtaagggtaaccggtAACgAATCTTGTCAACATGGCC | Rv HR1 BdCDPK4 | Gibson to 5’HA |
| BE_514 | gaaatcacatgatctgtcgacAAATTTTTCAATAAATGGCAATAGCTTATCAC | Fw HR2 BdCDPK4 | Gibson to glmS 3’ |
| BE_515 | ctcactatagaattcttaattaaGTGTAGATTTTGTCTTTGCCTATCG | Rv HR2 BdCDPK4 | Gibson-PacI |
| BE_518 | tgggatatattagccgtatcctaggGCGACTATAAAGCTGACAGACTTTGG | Fw HR1 BdCDPK7 | Gibson- AvrII |
| BE_519 | acatcgtaagggtaaccggtCTCCTCAACaGTGCCCAG | Rv HR1 BdCDPK7 | Gibson to 5’HA |
| BE_520 | gaaatcacatgatctgtcgacATGCCTAAGCGCCGTCG | Fw HR2 BdCDPK7 | Gibson to glmS 3’ |
| BE_521 | ctcactatagaattcttaattaaGTGACAGAGATTCAAGCTTGTACAGG | Rv HR2 BdCDPK7 | Gibson-PacI |
| BE_524 | tgggatatattagccgtatcctaggGCACATACACTCTATGTGGTACTCC | Fw HR1 BdPKAc1 | Gibson- AvrII |
| BE_525 | acatcgtaagggtaaccggtCCAGTTgTCaAAGGGGTCGG | Rv HR1 BdPKAc1 | Gibson to 5’HA |
| BE_526 | gaaatcacatgatctgtcgacTACTGTGAAAGTTCACGCAGAATTCC | Fw HR2 BdPKAc1 | Gibson to glmS 3’ |
| BE_527 | ctcactatagaattcttaattaaTGCATGAGTTGTTCAGAGTCAAACG | Rv HR2 BdPKAc1 | Gibson-PacI |
| BE_530 | tgggatatattagccgtatcctaggAACGCACGATTACCTTGCTCC | Fw HR1 BdPKAc2 | Gibson- AvrII |
| BE_531 | acatcgtaagggtaaccggtAACGTAAACGGCTCTGAGAAACG | Rv HR1 BdPKAc2 | Gibson to 5’HA |
| BE_532 | gaaatcacatgatctgtcgacCGGCATCATATTTTGCCTGAAATTTGG | Fw HR2 BdPKAc2 | Gibson to glmS 3’ |
| BE_533 | ctcactatagaattcttaattaaCCTGCACGGGAAAATGTCGC | Rv HR2 BdPKAc2 | Gibson-PacI |
| BE_536 | ACCGGTTACCCTTACGATGTTCCTGACTATGC | Fw 1xHA (Bd) for Knockdown tags. Universal for all. | Gibson to HR1 PCR (has AgeI) |
| BE_537 | gtcgacAGATCATGTGATTTCTCTTTGTTCAAGG | Rv glmS (Bd) for Knockdown tags. Universal for all. | Gibson to HR2 PCR (has SalI) |
| BE_551 | **CAAGTATCGATACAGGGAGGTT**GACAGTGCGATCATTGTTTG**gttttagagctaGAAAtagcaagttaaaataagg** | PLP1 guide PCR (or Gibson ssODN) |  |
| BE_552 | **CAAGTATCGATACAGGGAGGTT**TGTTTTAATGTTGTTCGTAT**gttttagagctaGAAAtagcaagttaaaataagg** | PLP3 guide PCR (or Gibson ssODN) |  |
| BE_553 | **CAAGTATCGATACAGGGAGGTT**CAAGGCCATGTTGACAAGAT**gttttagagctaGAAAtagcaagttaaaataagg** | CDPK4 guide PCR (or Gibson ssODN) |  |
| BE_554 | **CAAGTATCGATACAGGGAGGTT**TTGAATGAGTTAGTCTAATG**gttttagagctaGAAAtagcaagttaaaataagg** | CDPK5 guide PCR (or Gibson ssODN) |  |
| BE_555 | **CAAGTATCGATACAGGGAGGTT**GCTTAGGCATCACTCCTCAA**gttttagagctaGAAAtagcaagttaaaataagg** | CDPK7 guide PCR (or Gibson ssODN) |  |
| BE_556 | **CAAGTATCGATACAGGGAGGTT**ACAGTATCACCAGTTATCGA**gttttagagctaGAAAtagcaagttaaaataagg** | PKAc1 guide PCR (or Gibson ssODN) |  |
| BE_557 | **CAAGTATCGATACAGGGAGGTT**TCAGAGCCGTTTACGTTTAG**gttttagagctaGAAAtagcaagttaaaataagg** | PKAc2 guide PCR (or Gibson ssODN) |  |
| BE_563 | **CAAGTATCGATACAGGGAGGTT**GAGTCTATTTTGCATTATTC**gttttagagctaGAAAtagcaagttaaaataagg** | Guide Asp2 |  |
| BE_564 | tgggatatattagccgtatcctaggTTCGGATGATGCCAGTTGTCG | Fw HR1 Asp2 |  |
| BE_565 | acatcgtaagggtaaccggtTTTTGCATTATTCCGGTGCTTGG | Rv HR1 Asp2 |  |
| BE_566 | gaaatcacatgatctgtcgacACTCTTAATTTCAACATGTCGTGCG | Fw HR2 Asp2 |  |
| BE_567 | ctcactatagaattcttaattaaGGCAACCACTTCAAAGTTTGC | Rv HR2 Asp2 |  |
| BE_568 | **CAAGTATCGATACAGGGAGGTT**TATGCATCCATAGGATCATA**gttttagagctaGAAAtagcaagttaaaataagg** | Guide Asp3 |  |
| BE_569 | tgggatatattagccgtatcctaggAGTTGTTCGCGAGCACTATTGG | Fw HR1 Asp3 |  |
| BE_570 | acatcgtaagggtaaccggtGTAGTGTCTTGACGATGCAATCC | Rv HR1 Asp3 |  |
| BE_571 | gaaatcacatgatctgtcgacTGTATATATACtATATGATCCTATGGATGCATAC | Fw HR2 Asp3 |  |
| BE_572 | ctcactatagaattcttaattaaACAGCTACATGAGTCAGTAGAAAGG | Rv HR2 Asp3 |  |
| BE_573 | **CAAGTATCGATACAGGGAGGTT**TAATTAGTTATGTTTTAATA**gttttagagctaGAAAtagcaagttaaaataagg** | Guide DPAP1 |  |
| BE_574 | tgggatatattagccgtatcctaggCATACCTAACAGGAAAACGCATGC | Fw HR1 DPAP1 |  |
| BE_575 | acatcgtaagggtaaccggtTGTTTTAATATGGTTGTGTACATCGTGTAG | Rv HR1 DPAP1 |  |
| BE_576 | gaaatcacatgatctgtcgacCTAATTAATCGTGTTTTAAAATGCAACACGAC | Fw HR2 DPAP1 |  |
| BE_577 | ctcactatagaattcttaattaaGTAATGCTCCTTTATCTCCTTCCACATG | Rv HR2 DPAP1 |  |
| BE_589 SEQ | ATTTTGCATCCCCACGGTGG | Fw Bd U6 SEQ into guide |  |
| BE_590  SEQ | gtcacgacgttgtaaaacgacg | Rv Bd Cas9 plasmid SEQ into HR (constant on plasmid) |  |
| BE_596 | ACCTGAAACTCACGGACTTCG | Fw test PKAc1 tag Integration |  |
| BE_597 | CGAACCACTCCCGTATATGTCC | Rv test PKAc1 tag Integration |  |
| BE_598 | ACTTTGGTTTTGCCAAGCACG | Fw test PKAc2 tag Integration |  |
| BE_599 | GATGGCTGGTCATACAGTGC | Rv test PKAc2 tag Integration |  |
| BE_600 | GGAAAGGAAAAGTGTGTGGCG | Fw test PLP1 tag Integration |  |
| BE_601 | TCGCCTCAAACGCGGATTCC | Rv test PLP1 tag Integration |  |
| BE_602 | AAGACAAACATgtaaggagttgtgc | Fw test PLP3 tag Integration |  |
| BE_603 | TCTCGTATGCCAGTATGAAGTGC | Rv test PLP3 tag Integration |  |
| BE_604 | CATTATGTTTGGAGGTGCTGATGC | Fw test Asp2 tag Integration |  |
| BE_605 | CACAACAAGTCTCAAGATGAATAGGAGC | Rv test Asp2 tag Integration |  |
| BE_606 | GTTTGGTGGAGTGGATCCAAGG | Fw test Asp3 tag Integration |  |
| BE_607 | AGGTTTCATGAAGCTTTTGGTTTCG | Rv test Asp3 tag Integration |  |
| BE_608 | ACCTGCGATATGTTCAACCAGG | Fw test DPAP1 tag Integration |  |
| BE_609 | GATTTACTTCTTCTGCACGAGTGC | Rv test DPAP1 tag Integration |  |
| BE_610 | CATCAAACAGGTGCTCAGTGG | Fw test CDPK4 tag Integration |  |
| BE_611 | CAAAAGGCTGCATCTGGTATGC | Rv test CDPK4 tag Integration |  |
| BE_612 | GTAATGCGGCAGATCTTTTCTGC | Fw test CDPK5 tag Integration |  |
| BE_613 | GAACCACTTCCAAACCCCTGG | Rv test CDPK5 tag Integration |  |
| BE_614 | TCGTGAATTTTACGTGGCTTCTAGG | Fw test CDPK7 tag Integration |  |
| BE_615 | ATCAGTTGGTCAACACGTGGC | Rv test CDPK7 tag Integration |  |
| BE_550 | CCTAGGATACGGCTAATATATCCCATC | Rv Bd U6 terminator – for universal use with guide PCR |  |
| Synthesis 1 | aattctctagaGTGGTGTTAATTCATCGCTAGTAACCATATACGAGGGCATAGACAAGTCCTTCCACACGTGCATAGCCGCATTGGCTAGTAAGCTAAATATAGAGCAGATTTATGTCACCTTCAATTATTCTTTCTCAGCATGAATGTCACCTGTTTATGTCATAGTACGGCATAAACAGGCACGGCAATGGCACGGTCACCTTCTACATCACCAAACTACAATTGAAGTACAACAACAAAGTCAACTAATGAAGGTCTCCTCTATATCAAAGAGCGAGTCGTGTAGAGATAGTGATCCTCCGAAACTATTAAATGAAACCAGATATAGTGAAATGCTGTCCAACGTGCAGAAATAACCTTGCTTCATGATGTTTGGCAGCTCCCATATAGATCTGTACCATTTCAAACATTTTTATATGGCCATTTTTCTATTATATGTCGCATTGATCCTTATCCTTAATTTTGCATCCCCACGGTGGTACCTGTTGAGTGAGAGTCCCACTCCCCACAAGTATCGATACAGGGAGGTTggGTCTTCgaGAAGACctgttttagagctaGAAAtagcaagttaaaataaggctagtccgttatcaacttgaaaaagtggcaccgagtcggtgcTTTTTTTTTTGCATCTTTGCACAGAATATATCAATGATGGGATATATTAGCCGTATCCTAGGTATAGGAACTTTCGGGACCGTTAAATTGGTAGAGCATGAGGCCACAGGTGTAAGATTTGCTTTGAAGTGTGTTAGTAGGAAATGTATCCGTGCACTCAAGCAAGAGAAGCATATCAAGTTGGAAAGAGAAATAATGGCTCAGAATGACCATCCATTCATCGTTCAACTAGGTAATTGGTGCCGTGCCTTAAGCTAAATACTCTGCAGTAAAAACATTcAAaGAcGCTGATAATGTCTACTTtCTAcaAGAACTaGTGACTGGAGGTGAATTGTACGATGCAATCAGAAAAATTGGGTTGCTTTCGAGGTCTCAAGCACAATTTTACATAGCATCTATCGTCCTAGCATTCGAGTATTTGCATGAAAGACAAATCGCATATCGAGTAGGTTCAGAAGCTAAATGATTAATGCTTTCATAGGATTTGAAACCTGAAAATATACTTCTCGATGAACAGGGCTATATCAAACTGATTGATTTCGGATGTGCAAAAAAAATTAAAGAGGGCCTTAATTAA | The U6 promoter, bbs1 sites (for the guide), guide tracer/scafold, U6 terminator and PKG-T651Q repair template were synthesized by IDT |  |
